## Supplementary Information for "Quantifying accuracy and heterogeneity in single-molecule super-resolution microscopy"

#### This PDF file includes:

Supplementary Notes

**1 Supplementary Note 1: Deriving WIF based on Wasserstein gradient flows** **3**

**2 Supplementary Note 2: Landscape of the negative expected log likelihood near a saddle point** **6**

**3 Supplementary Note 3: Computing WIF** **8**

**5 Supplementary Note 5: Background estimation** **17**

**6 Supplementary Note 6: Localization confidence of an isolated molecule** **18**

**7 Supplementary Note 7: Effect of SNR on the performance of WIF in detecting position and brightness inaccuracies** **20**

**8 Supplementary Note 8: Quantifying localization accuracy via $\mathrm{WIF}_{\mathrm{avg}}$** **22**

**9 Supplementary Note 9: Pupil fitting** **23**

Supplementary Figures

S1 Landscape of expected, negative Poisson log likelihood in localizing two closely-spaced molecules 24
S2 Computing Wasserstein-induced flux (WIF) . . . 25
S3 Effect of PSF approximation on WIF as a function of the molecule's position relative to the computational grid and camera pixel size. . . . 26
S4 2D localization confidence of a molecule with various brightnesses . . . 27
S5 3D localization confidence of a molecule with various brightnesses . . . 28
S6 2D localization confidence of a molecule with various defocus mismatches . . . 29
S7 2D localization confidence of a molecule with various defocus and dipole-induced mismatches . . 30
S8 2D localization confidence of a bright molecule with various defocus and dipole-induced mismatches 31
S9 3D localization confidence and axial localization error of a molecule in a medium of mismatched refractive index . . . 32
S10 Effect of SNR on the performance of WIF in quantifying position inaccuracy . . . 33

S11 Effect of SNR on performance of WIF in quantifying brightness errors due to aberrated PSFs . . 34
S12 Effect of regularizer strength on WIF for accurate and inaccurate localizations . . . . . . . . . . 35
S13 Stability of WIF over a range of regularizer strengths for accurate and inaccurate localizations as a function of SNR . . . . . . . . . . . . . . . . . . . . . . . . . . . . . . . . . . . . . . . . . . . . . 36
S14 Using median and standard deviation of WIFs calculated over a range of regularizer strengths significantly improves detection of localization inaccuracies . . . . . . . . . . . . . . . . . . . . . . 37
S15 WIF confidences for two closely-located molecules at various separation distances . . . . . . . . . 38
S16 Wasserstein-induced flux ($\mathrm{WIF}_{\mathrm{avg}}$) quantifies localization accuracy without ground truth . . . . . 39
S17 Wasserstein-induced flux ($\mathrm{WIF}_{\mathrm{avg}}$) quantifies 3D localization accuracy without ground truth . . . 40
S18 WIF confidence map reveals artifacts in recovering a tubulin network from high-density SMLM data 41
S19 Localization confidences using uncalibrated and calibrated models for the 2D microtubule dataset 42
S20 3D model of the experimental Double-Helix PSF . . . . . . . . . . . . . . . . . . . . . . . . . . . 43
S21 WIF quantifies the accuracy of various 3D PSF models . . . . . . . . . . . . . . . . . . . . . . . . . 44
S22 WIF reveals sub-optimal fitting algorithms directly from experimental 3D SMLM data . . . . . . 45
S23 Comparison between Easy-DHPSF and RoSE in reconstructing the axial cross-section of a microtubule 46
S24 Validation of WIF on 3D experimental images of Alexa Fluor 647-labeled microtubules . . . . . . 47
S25 Filtering unreliable localizations using PSF width versus WIF for enhancing reconstruction accuracy 48
S26 Filtering unreliable localizations using localization precision versus WIF for enhancing reconstruction accuracy . . . . . . . . . . . . . . . . . . . . . . . . . . . . . . . . . . . . . . . . . . . . . 49
S27 Examples of heterogeneity in Nile red interactions with amyloid fibrils . . . . . . . . . . . . . . . . 50
S28 Execution time of computing WIF for various densities and two image sizes . . . . . . . . . . . . 51
S29 Effect of different measurement noise models on the accuracy of WIF . . . . . . . . . . . . . . . . 52
S30 Tuning the regularizer for optimal performance of WIF . . . . . . . . . . . . . . . . . . . . . . . . . 53

**Supplementary Tables**

S1 Mathematical notations used in RoSE . . . . . . . . . . . . . . . . . . . . . . . . . . . . . . . . . . 15
S2 Conditions and parameters used in computing WIF . . . . . . . . . . . . . . . . . . . . . . . . . . . 54

**Supplementary References**

### 1. Supplementary Note 1: Deriving WIF based on Wasserstein gradient flows

To gain insight into the origin of the localization uncertainty, let us formulate the localization task as minimizing the negative log-likelihood of observing an unknown number of molecules, $N$, each with a photon count $s_i$ and a position $\boldsymbol{r}_i$. If we know $N$ and assuming an imaging model with no mismatch, then the localization task reduces to simultaneously fitting $\{s_1, \boldsymbol{r}_1, \ldots, s_N, \boldsymbol{r}_N\}$ parameterizing our model to the observed data. The difficulty stems from not knowing $N$ *a priori*, which may cause localizations being practically trapped in a saddle point of the the landscape of the negative log-likelihood, while correct localizations correspond to *global minima* of this landscape. In our example of localizing two closely-spaced molecules (Fig. 1c,ii), almost all position estimates lie near $(0, 35\text{ nm})$, which exactly matches the centroid of the two true molecules. At the same time, the photon count estimates are twice as large as the ground-truth photons, and this point represents a saddle point of the negative log-likelihood (Fig. S1a). Similarly, model mismatches in the PSF may cause localizations to converge far from the global minima of the true negative log-likelihood.

A pivotal observation here is that these saddle points are unstable (in the sense of being a minimizer of the negative log-likelihood) upon a well-chosen perturbation (Supplementary Note 2). Put differently, for an accurate localization, the negative log-likelihood surface has a convex curvature as a function of the estimated position of a molecule. Therefore, if we locally perturb the position as well as photon count of a particular estimated molecule, relaxing this perturbation along the likelihood surface will most likely result in a localization very “close” to the unperturbed one. On the other hand, for an unreliable localization, we expect that the negative log-likelihood landscape changes arbitrarily in a local neighborhood (Fig. S1b). As a result, re-localizing most likely will *alter* the original localization. The stability in the position of a molecule upon a careful perturbation is precisely what we denote as the *quantitative confidence* of an SMLM localization. Motivated by this observation, we devise a robust method to measure the stability and therefore statistical confidence of *each localization* within an SMLM dataset.

**Localization stability for measuring confidence.** Intuitively, stability is a measure of discrepancy between a source point and a perturbed instance of this point after following a certain trajectory. To clarify, consider a strongly convex, differentiable function $f$ over some open set $\Omega \in \mathbb{R}$ taking its minimum at $\omega^* \in \Omega$. Since we are mostly interested in minimizers of some functional, as they are in a sense the best “fit” to the ground truth, we think of the confidence of a point estimate $\hat{\omega}$ as a measure of its distance to $\omega^*$. Since $\omega^*$ is unknown, we seek to measure the confidence of $\hat{\omega}$ without knowing $\omega^*$. To this end, we construct a simple single-step gradient-descent update and find a representation of stability to quantify the said confidence.

Consider the following gradient descent update given by the gradient-descent step with a small step size $\epsilon > 0$:

$$\omega_1 = \omega_0 - \epsilon \nabla f(\omega_0), \quad \omega_0 = \mathcal{P}(\hat{\omega}), \tag{1}$$

where $\omega_0$ is a local perturbation of $\hat{\omega}$ according to the operator $\mathcal{P}(\hat{\omega}) = \hat{\omega} + (1 - 2e)\Delta\hat{\omega}$ with $e \sim Bern(0.5)$ and perturbation distance $\Delta\hat{\omega} = |\hat{\omega} - \omega_0|$. Eq. (1) describes the movement of $\omega_0$ in the gradient vector field, $\nabla f$, transporting $\omega_0$ in the direction of decreasing $f$. If the estimate $\hat{\omega}$ is stable, we have $|\omega_1 - \hat{\omega}| < |\omega_0 - \hat{\omega}|$ as a result of our gradient-descent update, while for an unstable estimate, we can find a perturbation that results in $|\omega_1 - \hat{\omega}| > |\omega_0 - \hat{\omega}|$. Since $\omega^*$ is the minimizer of $f$, we have $|\omega_1 - \omega^*| < |\omega_0 - \omega^*|$ for *any* local perturbation of $\omega^*$. In other words, the gradient vector field pushes the perturbed point $\omega_0$ *toward* $\omega^*$. This observation tells us that we may quantify the confidence of $\hat{\omega}$ by measuring the average convergence of $\omega_0$ toward $\hat{\omega}$. We may define the confidence of a point $\hat{\omega}$ simply as

$$c = \frac{\mathbb{E}\left\{\text{sgn}\left[(\hat{\omega}-\omega_0)\cdot(\omega_1-\omega_0)\right]\cdot|\epsilon\nabla f(\omega_0)|\right\}}{\mathbb{E}\left[|\epsilon\nabla f(\omega_0)|\right]}, \tag{2}$$

where $\mathbb{E}$ denotes expectation over random perturbations and $\text{sgn}(x)$ takes the sign of a real number $x$. We call $c$ in Eq. (2) the normalized gradient flux, for reasons that become apparent later. A stable point has the maximum inward gradient flux, i.e., $c = 1$, while an unstable point has some degree of outward gradient flux, i.e., $c < 1$. Thus, $c$ represents a confidence score for any point in $\Omega$ without knowing $\omega^*$. As an example, for $f(\omega) = \omega^2$ thus implying $\omega^* = 0$, we find $c = \frac{2\Delta\hat{\omega}}{|\hat{\omega}-\Delta\hat{\omega}|+|\hat{\omega}+\Delta\hat{\omega}|}$. Obviously, $\hat{\omega} = \omega^* = 0$ is the most stable point with highest confidence, and the further away $\hat{\omega}$ is from 0, the worse the confidence.

We can gain more insight if we consider the recursive variational form of Eq. (1) as

$$\omega_k = \underset{\omega\in\Omega}{\arg\min}\left\{\frac{1}{2}\|\omega-\omega_{k-1}\|_2^2 + \epsilon_k f(\omega)\right\}, \quad k > 0. \tag{3}$$

Informally, Eq. (3) defines a discrete trajectory $\{\omega_k\}$ by minimizing $f$ while preserving a "local Euclidean distance" constraint. In the limit of $\epsilon_k \to 0$, i.e., considering continuous trajectories, we recover the *Cauchy Problem*, that is, $\frac{d\omega(t)}{dt} = -\nabla f(\omega(t))$, which defines the evolution of $\omega \in \Omega$ from an initial point $\omega_0$. The resulting curve $\{\omega(t)\}_{t\geq 0}$ is called a *gradient flow.*

**Wasserstein-induced flux.** Molecular brightnesses $s_i > 0$ and positions $\boldsymbol{r}_i \in \mathbb{R}^2$ in a single SMLM frame are expressed as $\mathcal{M} = \sum_{i=1}^N s_i\delta(\boldsymbol{r}-\boldsymbol{r}_i)$, which is a multi-parameter *distribution* in the space of non-negative finite measures $\mathrm{M}(\mathbb{R}^2)$. To extend our discussion to SMLM, we must define the distance between two candidate "guesses" $\mathcal{S}$ and $\mathcal{Q} \in \mathrm{M}(\mathbb{R}^2)$ for molecular parameters. We utilize the elegant theory of optimal transport, where roughly speaking, the optimal transport distance between any two measures is the minimum cost of transporting mass from one to the other as measured via some ground metric (1). The Wasserstein distance is particularly suitable, because its ground metric is simply Euclidean distance. The type-2 Wasserstein distance between two measures $\mathcal{S}, \mathcal{Q} \in \mathrm{M}(\mathbb{R}^2)$ is defined as

$$\mathbb{W}_2(\mathcal{S},\mathcal{Q}) = \min_{\pi\in\Pi(\mathcal{S},\mathcal{Q})}\sqrt{\int_{\mathbb{R}^2\times\mathbb{R}^2}\|\boldsymbol{r}-\boldsymbol{r}'\|_2^2\, d\pi(\boldsymbol{r},\boldsymbol{r}')}, \tag{4}$$

where $\Pi(\mathcal{S},\mathcal{Q})$ is the set of all couplings or *transportation plans* between $\mathcal{S}$ and $\mathcal{Q}$ satisfying a mass-conservation constraint (1). Equipped with Wasserstein distance, let us re-write the recursive dynamics of Eq. (3) as

$$\mathcal{S}_k = \underset{\mathcal{S}\in\mathrm{M}(\mathbb{R}^2)}{\arg\min}\left\{\frac{1}{2}\mathbb{W}_2^2(\mathcal{S},\mathcal{S}_{k-1}) + \epsilon_k\mathcal{L}(\mathcal{S})\right\}, \quad k > 0, \tag{5}$$

where $\mathcal{L} : \mathrm{M}(\mathbb{R}^2) \to \mathbb{R}$ is the negative Poisson log-likelihood, which is a convex functional.

We recall that our goal is to obtain a useful representation of point stability in the space of measures. Since stability is coupled with evolution of the measures, we analyze the properties of Wasserstein gradient flows, i.e., $\{\mathcal{S}\}_{t\geq 0}$. A set of intriguing results from the theory of Wasserstein gradient flow assert that if $\mathcal{S}$ has a smooth density, 1) there exists a unique transport map, $T_k : \mathrm{M}(\mathbb{R}^2) \to \mathrm{M}(\mathbb{R}^2)$, such that $\mathbb{W}_2^2(\mathcal{S},\mathcal{S}_{k-1}) = \int_\Omega |T_k(\boldsymbol{r})-\boldsymbol{r}|^2\, d\mathcal{S}_{k-1}(\boldsymbol{r})$; that is, the mass-weighted displacement distance for the transport plan is given by the type-2 Wasserstein distance; and 2) the *backward* velocity field $\mathbf{v}(\boldsymbol{r})$, i.e., the ratio between the displacement $T(\boldsymbol{r})-\boldsymbol{r}$ and the time step $\epsilon_k$, obtained in the transport of $\mathcal{S}_k$ to $\mathcal{S}_{k-1}$ is given by $\nabla\big(\frac{\delta\mathcal{L}}{\delta\mathcal{S}}(\mathcal{S})\big)(\boldsymbol{r})$ (in the limit of $\epsilon_k \to 0$) (2). The functional $\frac{\delta\mathcal{L}}{\delta\mathcal{S}}(\mathcal{S})$ is the so-called *first variation* of $\mathcal{L}$. In fact, the gradient flows satisfy the *continuity equation* (2):

$$\frac{\partial\mathcal{S}}{\partial t} - \nabla \cdot \Big(\mathcal{S}\nabla\big(\frac{\delta\mathcal{L}}{\delta\mathcal{S}}(\mathcal{S})\big)\Big) = 0. \qquad [6]$$

We now invoke the *divergence theorem* and define the *Wasserstein-induced flux* (WIF) corresponding to the perturbation volume around a molecule $\mathcal{V}$ as:

$$\begin{aligned}\text{WIF} &\triangleq \int_{\mathcal{V}} \bigg(\nabla \cdot \Big(\mathcal{S}\nabla\big(\frac{\delta\mathcal{L}}{\delta\mathcal{S}}(\mathcal{S})\big)\Big)\bigg) d\mathcal{V} \\ &= \int_{\mathfrak{S}} \Big(\mathcal{S}\nabla\big(\frac{\delta\mathcal{L}}{\delta\mathcal{S}}(\mathcal{S})\big) \cdot \boldsymbol{n}\Big)\, d\mathfrak{S},\end{aligned} \qquad [7]$$

where $\mathfrak{S}$ and $\boldsymbol{n}$ represent the closed surface on the boundary of $\mathcal{V}$ and its normal vector, respectively. We posit that WIF serves as a mathematically-grounded representation for stability that accounts for local interactions of point sources on the likelihood surface. We note source molecules have various brightnesses; photons are the conserved mass under our perturbation. We therefore normalize WIF w.r.t. the flux associated with an isolated source in $\mathcal{V}$. Henceforth, we denote WIF as its normalized quantity, which means that it takes on values within $[-1, 1]$ with 1 representing the maximum statistical confidence.

As stated previously, we can write the *gradient field* $\nabla\big(\frac{\delta\mathcal{L}}{\delta\mathcal{S}}(\mathcal{S})\big)(\boldsymbol{r}) = \mathbf{v}(\boldsymbol{r}) \approx [T_1(\boldsymbol{r}) - \boldsymbol{r}]/\epsilon$ in transporting $\mathcal{S}_1$ to $\mathcal{S}_0$. This equivalence effectively gives us a strategy to approximate WIF by finding an estimate of $T_1$, that is, the transport map. Unfortunately, it is computationally expensive to solve for the infinite-dimensional measure $\mathcal{S}_1$. In addition, molecules are in actuality point sources, which means that our object space $\mathrm{M}(\mathbb{R}^2)$ consists of discrete measures and not smooth densities. Even though the uniqueness condition of the transport map requires measures with smooth densities, we show that even with these approximations, our WIF dynamics mirror those predicted by Eq. (6). We designed an efficient, iterative algorithm to approximately compute $T_1$, which ultimately allows us to compute WIF (Supplementary Note 3, Fig. S2) using 1) raw SMLM images of blinking molecules and 2) a computational model of the imaging system.

### 2. Supplementary Note 2: Landscape of the negative expected log likelihood near a saddle point

In this section, we show how the landscape of the negative log likelihood changes around a saddle point, which represents a sub-optimal point. For clarity of the discussion, consider the problem of localizing two closely-spaced molecules. Without loss of generality, let us assume that they are located at $(0, r_y^*)$ and $(0, -r_y^*)$ with equal photon counts. Assuming a Gaussian PSF model, a noisy realization of their image $\boldsymbol{g} \in \mathbb{R}^m$ is formed according to the Poisson distribution:

$$\mu_i^* \triangleq A\Big\{s \exp\Big(-\frac{(u_i)^2 + (v_i - r_y^*)^2}{2\sigma^2}\Big) + s \exp\Big(-\frac{(u_i)^2 + (v_i + r_y^*)^2}{2\sigma^2}\Big)\Big\} + b_i, \tag{8}$$

$$g_i \sim \text{Pois}(\mu_i^*), \quad i \in \{1, \dots, m\}, \tag{9}$$

where $(u_i, v_i)$ are image-space coordinates sampled by the pixels of the camera, $A$ is a known normalizing constant, $\sigma$ is the PSF width, $\boldsymbol{b} \in \mathbb{R}^m$ denotes background, $m$ is the number of pixels, and Pois represents the Poisson probability distribution.

We further assume that $\sigma$, $\boldsymbol{b}$, and $s$ are known a priori. Therefore, we can write down the negative Poisson log-likelihood model $\mathcal{L}$, parameterized by an algorithm's estimated molecular positions $(r_x, r_y)$, and its expectation $\mathcal{E}$ over many realizations as follows:

$$\mu_i \triangleq A\left\{s \exp\left(-\frac{(u_i - r_x)^2 + (v_i - r_y)^2}{2\sigma^2}\right) + s \exp\left(-\frac{(u_i + r_x)^2 + (v_i + r_y)^2}{2\sigma^2}\right)\right\} + b_i, \quad i \in \{1, \dots, m\}. \tag{10}$$

$$\mathcal{L}(r_x, r_y; \boldsymbol{g}) = \sum_{i=1}^{m} \{\mu_i - g_i \log(\mu_i)\}. \tag{11}$$

$$\mathcal{E}(r_x, r_y) \triangleq \mathbb{E}\mathcal{L} = \sum_{i=1}^{m} \mu_i - \mathbb{E}\left(g_i \log(\mu_i)\right) = \sum_{i=1}^{m} \mu_i - \mu_i^* \log(\mu_i). \tag{12}$$

By taking the derivative of $\mathcal{E}$ with respect to $r_y$ we obtain

$$\frac{\partial \mathcal{E}}{\partial r_y} = \sum_{i=1}^{m} \frac{\partial \mu_i}{\partial r_y} - \frac{\mu_i^*}{\mu_i}\frac{\partial \mu_i}{\partial r_y} \quad \text{and} \tag{13}$$

$$\frac{\partial \mu_i}{\partial r_y} = \frac{As(u_i - r_y)}{\sigma^2} \exp\Big(-\frac{(u_i - r_x)^2 + (v_i - r_y)^2}{2\sigma^2}\Big) - \frac{As(u_i + r_y)}{\sigma^2} \exp\Big(-\frac{(u_i + r_x)^2 + (v_i + r_y)^2}{2\sigma^2}\Big). \tag{14}$$

A similar expression can be obtained for derivative of $\mathcal{E}$ w.r.t. $r_x$. It follows that $(r_x = 0, r_y = 0)$ is an equilibrium point of $\mathcal{E}$ and is located at center of $(0, r_y^*)$ and $(0, -r_y^*)$. In Fig. S1a, we plot the surface of $\mathcal{E}$ for various $(r_x, r_y)$ when $r_y^* = 35$ nm and $r_x^* = 0$. Interestingly, we observe that $(0, 0)$ is a saddle point.

As we discussed in the main text, when localizing two closely-spaced molecules, image-analysis algorithms often recover a single molecule whose position coincides with the saddle point of the negative log likelihood (Fig. 1c,ii). This phenomenon can be understood by noting that the algorithm does not know the number of underlying molecules, in this case two. This lack of knowledge can fool the algorithm to be trapped into a saddle point. To see this, we plot the surface of $\mathcal{E}$ parameterized only by one molecule (Fig. S1b). In this case, interestingly, $(0, 0)$ is an optimal point and the surface has an upward curvature around it. As can be seen from Fig. S1a, when we perturb the model by adding one more molecule, the seemingly optimal point $(0, 0)$ now corresponds to a saddle point, signaling its sub-optimality.

### 3. Supplementary Note 3: Computing WIF

Recall that our main goal is to compute an estimate of $T_1$, which characterizes the transport of mass (photons in our case) between the perturbed measure $\mathcal{S}_0$ and the solution to the following problem:

$$\mathcal{S}_1 = \underset{\mathcal{S}\in \mathrm{M}(\mathbb{R}^2)}{\arg\min} \left\{ \frac{1}{2}\mathbb{W}_2^2(\mathcal{S},\mathcal{S}_0) + \epsilon\mathcal{L}(\mathcal{S}) \right\}, \tag{15}$$

Given such an estimate, we then are able to compute the vector field $\mathcal{S}\nabla\left(\frac{\delta\mathcal{L}}{\delta\mathcal{S}}(\mathcal{S})\right)$ and, thus, WIF according to

$$\mathrm{WIF} = \int_{\mathfrak{S}} \left(\mathcal{S}\nabla\left(\frac{\delta\mathcal{L}}{\delta\mathcal{S}}(\mathcal{S})\right)\cdot \boldsymbol{n}\right)\, d\mathfrak{S}, \tag{16}$$

where $\mathfrak{S}$ represents the closed surface on the boundary of a chosen perturbation volume $\mathcal{V}$ and $\boldsymbol{n}$ is the vector normal to $\mathfrak{S}$. As stated in the main text, we consider discrete measures, i.e., point sources, to obtain a discrete version of WIF in Eq. (7). As a first task, we start by showing how to perturb a set of localizations or point sources.

To proceed, we discretize the underlying object space, $\mathbb{R}^2$, into a square grid of $\mathcal{N}$ points separated by $2\rho$ (Fig. S2, Table S2). We denote $\mathcal{G}$ and $\{r_{\mathcal{G}_i}\}_{1:\mathcal{N}}$ as the grid and its points, respectively. Assuming that any two point sources are separated by at least $\rho$, we then apply a reparameterization trick such that any *discrete* measure $\mathcal{M} = \sum_{i=1}^{N} s_i\delta(\boldsymbol{r}-\boldsymbol{r}_i)$, that is, a collection of $N$ point sources located at $\{\boldsymbol{r}_1,\ldots,\boldsymbol{r}_N\}$ with brightness $\{s_1,\ldots,s_N\}$ , can be written as

$$\mathcal{M} = \sum_{i=1}^{N} s_{[i]}\delta\left(\boldsymbol{r} - (\boldsymbol{r}_{\mathcal{G}_{[i]}} + \Delta\boldsymbol{r}_{[i]})\right), \tag{17}$$

with grid point index $[i] \in \{1,\ldots,\mathcal{N}\}$ and the distance from the point source to the nearest grid point $\left\|\Delta\boldsymbol{r}_{[i]}\right\|_2 \le \rho$ (Fig. S2).

***Perturbing a set of localizations.*** Let $\hat{\mathcal{M}} = \sum_{i=1}^{\hat{N}} \hat{s}_i\delta(\boldsymbol{r}-\hat{\boldsymbol{r}}_i)$ be the localization estimates corresponding to $\hat{N}$ molecules fed to the confidence mapping algorithm. We can equivalently represent these estimates using our constructed grid as $\hat{\mathcal{M}} = \sum_{i=1}^{\hat{N}} \hat{s}_{[i]}\delta\left(\boldsymbol{r} - (\hat{\boldsymbol{r}}_{\mathcal{G}_{[i]}} + \Delta\hat{\boldsymbol{r}}_{[i]})\right)$. We assume that $\hat{s} = \sum_{i=1}^{\hat{N}} \hat{s}_{[i]}$ equals the total mass of (i.e., photons detected from) the ground-truth sources. The perturbed measure $\mathcal{M}_0$ is defined as

$$\mathcal{P}(\hat{\mathcal{M}}) = \mathcal{M}_0 \triangleq \sum_{i=1}^{\hat{N}} \left(\sum_{j=1}^{8} \hat{s}_{[i,j]}\delta(\boldsymbol{r} - \hat{r}_{\mathcal{G}[i,j]})\right), \tag{18}$$

where $\hat{s}_{[i,j]} = \hat{s}_{[i]}/8$. Eq. (18) states that for each point source in $\hat{\mathcal{M}}$, we redistribute its photons to 8 point sources located at the closest neighboring grid points represented as $\{\hat{r}_{\mathcal{G}_{[i,1]}},\ldots,\hat{r}_{\mathcal{G}_{[i,8]}}\}$ (Fig. S2). For convenience, we index these points as $\mathrm{Nh}([i]) = \{[i,1],\ldots,[i,8]\}$. This perturbation $\mathcal{M}_0$ thus has symmetric distributions around the original point sources $\hat{\mathcal{M}}$.

As mentioned, we are interested in a unique map that describes the transport of mass between two measures. One way to achieve this is to directly regularize mass transportation. To this end, we consider a local constraint on $\mathcal{M}$ as

$$\mathcal{C} = \{(s_i,\Delta r_i) \mid \|\Delta r_i\|_2 \le \rho, i \in \mathrm{Supp}(\mathcal{M}_0)\}, \tag{19}$$

where the support $\mathrm{Supp}(\mathcal{M}_0)$ is defined as $\left\{j \in \{1, \cdots, \mathcal{N}\} \mid j \in \bigcup_{i=1}^{\hat{N}} \mathrm{Nh}([i])\right\}$. Effectively, this constraint forces each point to be transported along a unique trajectory in a local neighborhood of the unperturbed source. We propose to solve the following regularized one-step dynamical process:

$$\mathcal{M}_1 = \underset{\mathcal{M} \in \mathrm{M}(\mathbb{R}^2) \cap \mathcal{C}}{\arg\min} \left\{ \frac{1}{2} \mathbb{W}_2^2(\mathcal{M}, \mathcal{M}_0) + \epsilon \mathcal{L}(\mathcal{M}) \right\}. \tag{20}$$

In order to solve for Eq. (20) efficiently, we propose to bound $\mathbb{W}_2^2(\mathcal{M}, \mathcal{M}_0)$ from above with a group-sparsity norm. We will show that such a relaxation allows us to derive a convex program for approximating $\mathcal{M}_1$.

***Bounding the square of Wasserstein distance.*** Our goal is to show that $\forall \mathcal{M} \in \mathrm{M}(\mathbb{R}^2) \cap \mathcal{C}$ we can bound $\mathbb{W}_2^2(\mathcal{M}, \mathcal{M}_0)$ from above with

$$\sum_{i=1}^{\hat{N}} \sum_{j=1}^{8} \sqrt{s_{[i,j]}^2 + s_{[i,j]}^2 \left\| \Delta \boldsymbol{r}_{[i,j]} \right\|_2^2}, \tag{21}$$

so long as $\left\| \Delta \boldsymbol{r}_{[i,j]} \right\|_2^2 \leq \sqrt{1 + \left\| \Delta \boldsymbol{r}_{[i,j]} \right\|_2^2}$. We note that this assumption can be easily satisfied by appropriately scaling $2\rho$, the separation between grid points, in the object model (Fig. S2).

First, we notice that using our reparameterization trick in Eq. (17), any measure $\mathcal{M} \in \mathrm{M}(\mathbb{R}^2) \cap \mathcal{C}$ can be represented as $\mathcal{M} = \sum_{i=1}^{\hat{N}} \left( \sum_{j=1}^{8} s_{[i,j]} \delta \left( \boldsymbol{r} - (\boldsymbol{r}_{\mathcal{G}_{[i,j]}} + \Delta \boldsymbol{r}_{[i,j]}) \right) \right)$ such that $\sum_{i=1}^{\hat{N}} \sum_{j=1}^{8} s_{[i,j]} = \hat{s}$, that is, detected photons are preserved. Therefore,

$$\begin{aligned} \mathbb{W}_2^2(\mathcal{M}, \mathcal{M}_0) &\leq \sum_{i=1}^{\hat{N}} \sum_{j=1}^{8} s_{[i,j]} \left\| \Delta \boldsymbol{r}_{[i,j]} \right\|_2^2 && [22] \\ &\leq \sum_{i=1}^{\hat{N}} \sum_{j=1}^{8} s_{[i,j]} \sqrt{1 + \left\| \Delta \boldsymbol{r}_{[i,j]} \right\|_2^2} && [23] \\ &= \sum_{i=1}^{\hat{N}} \sum_{j=1}^{8} \sqrt{s_{[i,j]}^2 + s_{[i,j]}^2 \left\| \Delta \boldsymbol{r}_{[i,j]} \right\|_2^2}, && [24] \end{aligned}$$

where the first inequality in Eq. (22) follows from the definition of Wasserstein distance and the second inequality in Eq. (23) is the consequence of our assumption. Notice that Eq. (24) may be recast as a group-sparsity norm:

$$\sum_{i=1}^{\hat{N}} \sum_{j=1}^{8} \sqrt{s_{[i,j]}^2 + s_{[i,j]}^2 \left\| \Delta \boldsymbol{r}_{[i,j]} \right\|_2^2} = \sum_{i=1}^{\mathcal{N}} \sqrt{s_i^2 + s_i^2 \left\| \Delta \boldsymbol{r}_i \right\|_2^2} \triangleq \mathcal{R}(\mathcal{M}), \tag{25}$$

where it is assumed that $s_i = 0$ for grid index $i$ that do not contain perturbed sources, i.e., $\{s_i = 0 \mid i \notin \mathrm{Supp}(\mathcal{M}_0)\}$.

***Confidence quantification via convex programming.*** We re-write the objective objective function in Eq. (20), bounding $E(\mathcal{M})$ from above:

$$E(\mathcal{M}) = \frac{1}{2}\mathbb{W}_2^2(\mathcal{M}, \mathcal{M}_0) + \epsilon\mathcal{L}(\mathcal{M}) + \mathcal{I}_C(\mathcal{M}) \quad [26]$$

$$\leq \frac{1}{2}\mathcal{R}(\mathcal{M}) + \epsilon\mathcal{L}(\mathcal{M}) + \mathcal{I}_C(\mathcal{M}), \quad [27]$$

where $\mathcal{I}_\mathcal{C}$ represents the indicator function of the constraint set $\mathcal{C}$:

$$\mathcal{I}_\mathcal{C}(\mathcal{M}) = \begin{cases} 0, & \text{if} \quad \mathcal{M} \in \mathcal{C} \\ \infty, & \text{otherwise} \end{cases}. \quad [28]$$

Based on this observation, we consider the following regularized one-step dynamical process:

$$\mathcal{M}_1 = \underset{\mathcal{M}\in\mathrm{M}(\mathbb{R}^2)}{\arg\min} \left\{\frac{1}{2}\mathcal{R}(\mathcal{M}) + \epsilon\mathcal{L}(\mathcal{M}) + \mathcal{I}_C(\mathcal{M})\right\}. \quad [29]$$

Fortunately, Eq. (29) is a convex program that can be efficiently solved using optimization techniques developed in Ref. (3) and detailed in Ref. (4).

We observe that we can write $\mathcal{M}_1 = \sum_{i=1}^{\hat{N}} \left( \sum_{j=1}^{8} \tilde{s}_{[i,j]} \ \delta\left(\boldsymbol{r} - \boldsymbol{r}_{[i,j]}\right)\right)$ by our construction as:

$$\mathcal{M}_1 = \sum_{i=1}^{\hat{N}} \Big( \sum_{j=1}^{8} \tilde{s}_{[i,j]} \ \delta\left(\boldsymbol{r} - (\hat{\boldsymbol{r}}_{\mathcal{G}_{[i,j]}} + \Delta\tilde{\boldsymbol{r}}_{[i,j]})\right)\Big). \quad [30]$$

Comparing the expressions of $\mathcal{M}_1$ and $\mathcal{M}_0$ we deduce that, assuming $\hat{s}_{[i,j]} = \tilde{s}_{[i,j]}$, the displacement $T_{1[i,j]} - \boldsymbol{r}_{[i,j]}$ in transporting $\mathcal{M}_1$ to $\mathcal{M}_0$ is simply given by $-\Delta\tilde{r}_{[i,j]}$, the backward displacement vector at each grid point. Recall in calculating WIF as in Eq. (7), we may replace $\nabla\big(\frac{\delta\mathcal{L}}{\delta\mathcal{M}}(\mathcal{M})\big)(\boldsymbol{r})$ with $\mathbf{v}(\boldsymbol{r}) \approx \left[T_1(\boldsymbol{r}) - \boldsymbol{r}\right]/\epsilon$. Consequently, we can compute an approximate WIF, i.e., localization confidence, for the $i^{\text{th}}$ molecule as follows:

$$\mathrm{WIF}_i = c_i = \int_{\mathfrak{S}} \Big(\mathcal{S}\nabla\big(\frac{\delta\mathcal{L}}{\delta\mathcal{S}}(\mathcal{S})\big)\cdot\boldsymbol{n}\Big)\, d\mathfrak{S}, \quad [31]$$

$$\approx \frac{\sum_{j=1}^{8} \hat{s}_{[i,j]} \left\|\Delta\tilde{\boldsymbol{r}}_{[i,j]}\right\|_2 \cdot \cos(\zeta_{[i,j]})}{\sum_{j=1}^{8} \hat{s}_{[i,j]} \left\|\Delta\tilde{\boldsymbol{r}}_{[i,j]}\right\|_2}, \quad [32]$$

where

$$\cos(\zeta_{[i,j]}) = \frac{\Delta\tilde{\boldsymbol{r}}_{[i,j]}^{\mathrm{T}}(\hat{\boldsymbol{r}}_{[i]} - \hat{\boldsymbol{r}}_{\mathcal{G}_{[i,j]}})}{\left\|\Delta\tilde{\boldsymbol{r}}_{[i,j]}\right\|_2 \left\|\hat{\boldsymbol{r}}_{[i]} - \hat{\boldsymbol{r}}_{\mathcal{G}_{[i,j]}}\right\|_2}, \quad [33]$$

$\hat{\boldsymbol{r}}_{[i]}$ is the original estimated position of $i^{\text{th}}$ molecule, $i \in \{1, \ldots, \hat{N}\}$, and T denotes the transpose operator of a matrix. We call $\zeta_{[i,j]}$ the transport angle as it represents the angle between the estimated source molecule and displacement from $\hat{\boldsymbol{r}}_{\mathcal{G}_{[i,j]}}$ (Fig. S2).

**A. Extending WIF to 3D SMLM.** A natural extension of WIF to 3D imaging involves locally perturbing an estimated molecule within a small volume. Since the optical PSF is not shift-invariant along z as it is along x and y, such a strategy requires the PSF model to be computed individually for each molecule in the imaging volume, thereby complicating the computation. With this complexity in mind, we consider a variant of WIF in 3D that lends itself to an efficient algorithm, which is identical to that of WIF in 2D. Specifically, we capitalize on the observation that an accurate localization in 3D should not only be stable w.r.t. a volumetric perturbation but should also be stable w.r.t. perturbation within the xy plane. Therefore, for any 3D localization, WIF performs a 2D (in-plane) perturbation similar to Eq. 18, in which each perturbed source molecule maintains the same axial position of the original estimated molecule.

***Local perturbation in 3D.*** Similar to the 2D case, we perturb a set of localizations $\hat{\mathcal{M}}$ by introducing a small distortion in the positions and brightnesses of the molecules in $\hat{\mathcal{M}}$ to produce another set of localizations $\mathcal{M}_0$:

$$\mathcal{M}_0 = \sum_{i=1}^{\hat{N}} \sum_{j=1}^{8} \hat{s}_{[i,j]} \delta(\boldsymbol{r} - \hat{\boldsymbol{r}}_{[i,j]}), \tag{34}$$

where $\sum_{j=1}^{8} \hat{s}_{[i,j]} = \hat{s}_{[i]}$ and $\hat{\boldsymbol{r}}_{[i,j]}$ is one of the 8 neighboring grid points of $\hat{\boldsymbol{r}}_{\mathcal{G}_{[i]}}$ (Figure S2) such that $\hat{\boldsymbol{r}}_{[i,j]}|_z = \hat{\boldsymbol{r}}_i|_z$, that is, each perturbed molecule located at $\hat{\boldsymbol{r}}_{[i,j]}$ and the original molecule located at $\hat{\boldsymbol{r}}_i$ have the same axial position. We next solve a regularized transport problem (Eq. 29) assuming that the PSF in the negative log likelihood ($\mathcal{L}$) is evaluated at $z = \hat{\boldsymbol{r}}_i|_z$.

**B. Accounting for pixel-dependent readout noise for sCMOS cameras.** In contrast to EMCCD cameras, in which the readout noise can be effectively neglected due to large amplification gain, the readout noise in sCMOS cameras can be significant, especially in cases where the background is only a few photons per pixel (5). The probability distribution of photon counts in each pixel may be modeled as a convolution of Poisson shot noise with a Gaussian distribution due to readout noise. Therefore, a simplified Poisson noise model, which ignores the readout noise, may produce sub-optimal localizations. Fortunately, we can still approximate the convolved distribution using a shifted Poisson distribution as described below.

Recall that the expected number of photons detected at pixel $i$ is denoted by $\mu_i$. We further denote the variance of the readout noise at pixel $i$ by $var_i$. Then one can show that the following (approximately) holds:

$$\mu_i + var_i \sim \text{Poiss}(\mu_i + var_i), \tag{35}$$

where Poiss($\cdot$) denotes the Poisson distribution (5). We stress that WIF quantifies the accuracy of a localization algorithm and a computational model, which includes noise statistics. In particular, if in some applications the readout noise is significant, we may use an augmented formulation of WIF according to Eq. (35), which uses the Poisson negative log likelihood in its objective function.

An interesting and useful application of WIF could be to compare the accuracy of two localization algorithms that use different noise models. Specifically, we consider an imaging experiment where there exists a pixel-dependent readout noise, which models a typical sCMOS camera. We consider a camera whose pixels exhibit a Gaussian-distributed readout noise with a standard deviation of 1 photon, except for one pixel located near the center of the field-of-view that has a standard deviation of 20 photons (Fig. S29b). We generate 200 images of a SM (Fig. S29a) in which each realization is obtained by summing two images, one that is sampled from a Poisson distribution whose expected number of photons is given by the ideal microscope PSF and the other one sampled from a Gaussian distribution matching the camera's readout noise map. In each noisy image, we set all pixels with values smaller than or equal to

0 to 0.0001. We then use RoSE to localize the SM with the Poisson noise model (simplified), which ignores readout noise, as well as the sCMOS noise model in Eq. (35). As shown in Fig. S29, the localization precision and accuracy obtained using the sCMOS noise model is significantly better than those obtained via the Poisson noise model. We next compute the corresponding WIF scores using the sCMOS noise model, revealing that the simplified noise model produces localizations with an average WIF of only 0.65, while the sCMOS noise model has a WIF average of 0.75. These results demonstrate that WIF is able to quantify the superiority of a noise model.

However, if WIF algorithm uses the same noise model as the localization algorithm, albeit simplified one, the WIF algorithm may overfit and produces higher values for simplified noise model despite having worse precision and accuracy. To quantify this behavior, we use the simplified noise model in WIF algorithm to analyze the localizations in Fig. S29b. As shown in Fig. S29e, we see that the distribution of confidences for the simplified case is better (mean confidence 0.63 versus 0.51) but still worse than the distribution obtained using the sCMOS noise model.

**C. Parameters.** In computing WIF, we specify a grid $\mathcal{G}$ in object space that defines our perturbation $\mathcal{M}_0$ of the molecule parameters $\hat{\mathcal{M}}$ obtained from an SMLM algorithm, as described above (Fig. S2). We intuitively choose the grid spacing $2\rho$ to be comparable to the localization precision, as we are interested in quantifying some parameters related to the variations in the likelihood landscape. Additionally, since we are interested in detecting errors in high-density SM localization, we note that $\rho$ needs to be selected such that the perturbed sources associated with two closely-located molecules are separated. Fortunately, since practically it is not possible to localize two closely-located sources below ~100 nm for typical SMLM SNRs, our choice of $\rho$ (tens of nanometers) does not present a bottleneck. Finally, due to our implementation of WIF, we choose a small $\rho$ comparable to localization precision to avoid errors caused by PSF approximation (see Supplementary Note 3D and Fig. S3). We elect $2\rho =$ (image pixel size)$/k$ for some integer $k > 0$ (Table S2).

Another consideration is the choice of $\epsilon$ in Eq. (29). Note that we can rewrite Eq. (29) as:

$$\mathcal{M}_1 = \underset{\mathcal{M}\in \mathrm{M}(\mathbb{R}^2)}{\arg\min} \left\{ \nu\mathcal{R}(\mathcal{M}) + \mathcal{L}(\mathcal{M}) + \mathcal{I}_C(\mathcal{M}) \right\}, \qquad [36]$$

where we have defined $\nu = 1/(2\epsilon)$. In essence, $\nu$ relates to our degree of uncertainty in $\mathcal{M}_0$, i.e., the perturbed localizations. For large $\nu$, we have little uncertainty in $\mathcal{M}_0$ and the solution to Eq. (36) is simply $\mathcal{M}_0$. On the other hand, if we expect that $\mathcal{M}_0$ is uncertain, or equivalently, the unperturbed localizations $\hat{\mathcal{M}}$ are uncertain, we may choose a small value for $\nu$. Our simulation results show that when localizations are accurate, WIF is not so sensitive with respect to the choice of $\nu$ (Fig. S12a). However, for inaccurate localizations WIF can be sensitive to $\nu$ (Fig. S12b). As mentioned in the main text, such a property can be exploited to obtain a degree of stability for WIF, which improves the detection performance of WIF. In particular, we can compute WIFs at various regularizer strengths and compute median absolute deviation (MAD) and median statistics. The drawback of this methods is that it may not be computationally efficient. Therefore, we may tune $\nu$ based on some training data.

In this paper, we use isolated images of molecules generated from a vectorial image-formation model or obtained via control experiments (Methods) as training data. We used a simple tuning method as depicted in Fig. S30 for optimizing the regularizer strength. Concretely, we generate 500 images of an isolated molecule with expected brightness and background according to the experimental conditions. Next, we localize these molecules using RoSE. Next, we add position errors, which are randomly selected within $[33, 66]$ nm, to localizations obtained by RoSE. We denote these corrupted localizations as inaccurate ones while call those obtained by RoSE as accurate ones. Next, we feed both types of localizations, accurate and inaccurate, to the WIF algorithm using various regularizer strengths. Using a WIF threshold of 0.5, we classify the localizations with WIF greater than 0.5 as accurate and those with WIF less than 0.5 as inaccurate. Based on these classified localizations, we compute the Jaccard indices for each regularizer strength. Finally, we select the regularizer strength with the maximum Jaccard index as the optimal one. Note that we have found that 0.1 for $\nu$ generally works well for various experimental conditions. We list all the parameters used for computing WIF in Table (S2).

**D. PSF model.** In order to efficiently solve the optimization problem in Eq. (36), we use a first-order approximation of the exact PSF, as in Ref. (3). Recall that for any $\mathcal{M} \in \mathrm{M}(\mathbb{R}^2)$ we have:

$$\mathcal{M} = \sum_{i=1}^{\mathcal{N}} s_{[i]} \delta \left( \boldsymbol{r} - (\boldsymbol{r}_{\mathcal{G}_{[i]}} + \Delta \boldsymbol{r}_{[i]}) \right), \tag{37}$$

where $\left\| \Delta \boldsymbol{r}_{[i]} \right\|_2 \le \rho$. Given these molecular parameters $\mathcal{M} \in \mathrm{M}(\mathbb{R}^2)$ and the integrated PSF $q_{c,j}(\boldsymbol{r})$ of a molecule located at position $\boldsymbol{r}$ (see Methods), the resulting intensity $\mu_j$, that is, the expected number of photons detected in camera pixel $j \in \{1, \ldots, m\}$, can be written as

$$\mu_j = \sum_{i=1}^{\mathcal{N}} s_{[i]} q_{c,j} \left( \boldsymbol{r}_{\mathcal{G}_{[i]}} + \Delta \boldsymbol{r}_{[i]} \right) + b_j \tag{38}$$

$$= \sum_{i=1}^{\mathcal{N}} s_{[i]} q_{c,j} \left( \boldsymbol{r}_{\mathcal{G}_{[i]}} \right) + s_{[i]} q'_{c,j} \left( \boldsymbol{r}_{\mathcal{G}_{[i]}} \right) \Delta \boldsymbol{r}_{[i]} + s_{[i]} O \left( \Delta \boldsymbol{r}_{[i]}^2 \right) + b_j, \tag{39}$$

where $q'_{c,j} \left( \boldsymbol{r}_{\mathcal{G}_{[i]}} \right)$ denotes the derivative of $q_{c,j}$ with the respect to position evaluated at $\boldsymbol{r}_{\mathcal{G}_{[i]}}$ and $O \left( \Delta \boldsymbol{r}_{[i]}^2 \right)$ represents the residual error in the Taylor expansion of $q_{c,j}$. In our implementation, we drop the residual term in Eq. (39) and only consider terms up to first order:

$$\mu_j = \sum_{i=1}^{\mathcal{N}} s_{[i]} q_{c,j} \left( \boldsymbol{r}_{\mathcal{G}_{[i]}} \right) + s_{[i]} q'_{c,j} \left( \boldsymbol{r}_{\mathcal{G}_{[i]}} \right) \Delta \boldsymbol{r}_{[i]} + b_j. \tag{40}$$

Due to this model approximation, for molecules whose positions are within the grid, their confidences are slightly reduced, but still remain above 0.8 (Fig. S3a). We also investigated the effect of first-order approximation errors for a large camera pixel size (Fig. S3b). A larger camera pixel size (160 nm, median WIF of 0.84) introduces larger residual errors as compared to a smaller camera pixel size (58.5 nm, median WIF of 0.94), which adversely impact the estimated WIFs. Finally, note that Eq. (40) can be readily extended to 3D.

**E. Quantifying the stability of WIF.** We propose to quantify the stability/variance of a WIF estimate by computing WIF values at various regularizer strengths. Let $c_i(\nu_j)$ be the estimated WIF at the regularizer strength of $\nu_j$ (see Eq. 36). We consider a range of regularizer strengths $[0.06, 0.12]$ and compute WIFs for a set of $\nu_j$ (typically 10-13 values for $\nu$). We next compute the mean absolute deviations (MAD) of the estimated WIFs and obtain an estimate for the standard deviation of WIF as $1.48 \times$ MAD.

**F. Implementation and computational complexity.** The main computational task in computing WIF is solving Eq. (36). As detailed in Ref. (3), we can efficiently solve this convex optimization problem using accelerated proximal gradient algorithms, which are iterative methods. Therefore, the main computational cost is computing the gradients of $\mathcal{L}$ in Eq. (36). Assuming a square input image with size $\sqrt{m} \times \sqrt{m}$ pixels, it can be shown that such gradients can be computed with $O\left(m \log_2(m)\right)$ cost using the fast Fourier transform.

Here, we use a simple implementation of our algorithm from Ref. (3) in Matlab (R2018a) on a desktop computer with 20 GB RAM and an Intel® Core™ i7-6700 CPU (3.40 GHz clock). We plot the execution time of computing WIF for various densities and two image sizes in Fig. (S28). In the future, our implementation can be accelerated exploiting massive parallelism (e.g., segmenting a large field of view into smaller images and analyzing multiple frames simultaneously via graphical processing units).

### 4. Supplementary Note 4: Localization softwares

**ThunderSTORM** (6). For analyzing all datasets via the ThunderSTORM plugin in ImageJ (7), we used Wavelet filter (B-Spline) (scale= 2, order= 3), local maximum selection with a threshold of 1.2× std(Wave.F1), nmax = 3 (maximum number of molecules within the fitting area), and an integrated Gaussian PSF. We used fitradius= 4 (Figs. 1, 4, 5, S6, S7, S8) and fitradius= 3 (Figs. 2, S19). For Fig. S16, we enabled multi-emitter analysis with a p-value of $10^{-6}$.

**RoSE** (3). The regularizer parameter was empirically set to 0.19 (Figs. 1, S3, S4, S6, S7, S8, S10-S16), 0.3 (Figs. 2, S19), 0.08 (Figs. 3, S22g), 0.21 (Figs. 4, 5, S18, S19), and 0.15 (Figs. S5, S9, S17), 0.3 (Fig. S20), and 0.18 (Fig. S22b,c,f).

**FALCON** (8). We used the default settings of FALCON software (Figs. S16, S18).

**SQUIRREL** (9). We used default settings of the SQUIRREL plugin in ImageJ. The resolution scaling function (RSF), equivalent to the PSF, was estimated by SQUIRREL directly on the low-density blinking data (Fig. S18).

#### A. Optimizing fitting routines within RoSE.

Here, we provide a description of fitting routines within RoSE. In particular, we focus on second stage of this algorithm in which an adaptive, constrained maximum likelihood problem is solved. Let us define some notations for ease of discussion:

**Table S1. Mathematical notations used in RoSE**

| Notation | Definition |
|---|---|
| $N$ | number of grid points |
| $\boldsymbol{s} \in \mathbb{R}^N$ | brightnesses at grid points |
| $\Delta\boldsymbol{x} \in \mathbb{R}^N$ | position offsets along x at grid points |
| $\Delta\boldsymbol{y} \in \mathbb{R}^N$ | position offsets along y at grid points |
| $\Delta\boldsymbol{z} \in \mathbb{R}^N$ | position offsets along z at grid points |
| $\boldsymbol{\gamma} = [\boldsymbol{s}, \Delta\boldsymbol{x}, \Delta\boldsymbol{y}, \Delta\boldsymbol{z}] \in \mathbb{R}^{4N}$ | all parameters at grid points |
| $\gamma_{\text{init}}$ | an initial estimate |
| $\mathcal{L}(\boldsymbol{\gamma})$ | the negative log likelihood evaluated at $\boldsymbol{\gamma}$, |
| $\nabla\mathcal{L}_{\boldsymbol{\gamma}}(\boldsymbol{v})$ | the derivative of the negative log likelihood w.r.t. $\boldsymbol{\gamma}$ evaluated at $\boldsymbol{v}$, |
| Supp($\boldsymbol{\gamma}$) | the set of grid points that contain one molecule |
| DistSupp($\boldsymbol{\gamma}$) | computes the set of nearest neighbor grid points (i.e., have minimum distance) to each localized molecule |
| UpdateSupp($\boldsymbol{\gamma}$) | updates the support of $\boldsymbol{\gamma}$ so that every molecule is assigned to its closest grid point |
| $\beta \in \mathbb{R}_+$ | step size |
| $t$ | iteration number |
| $\eta > 1$ | backtracking parameter |

When localizing molecules within noisy images, an initial estimate $\boldsymbol{\gamma}_{\text{init}}$ may happen to be far from the true parameter, e.g., the position of a molecule. In these scenarios, an algorithm that constructs a local PSF model around current estimate should adaptively update this model (i.e., the object support), thereby enabling precise and accurate localization. In particular, RoSE builds local PSF models around grid points that contain a molecule and minimizes $\mathcal{L}(\cdot)$ in the neighborhood of each of those points. Therefore, an adaptive step is introduced in RoSE to adjust the current support Supp($\boldsymbol{\gamma}$) to match the closest grid points. Concretely, the fitting routines of RoSE can be described as follows:

**Algorithm 1** Solving an adaptive, constrained maximum likelihood problem within RoSE

**Input**: $\{\boldsymbol{\gamma}_{\text{init}}, \beta_{\text{init}}\}$

**Step** 0. Take $\boldsymbol{v}_1 = \boldsymbol{\gamma}_{\text{init}}, t_1 = 1,$ and $\eta > 1$. Set $\beta_0 = \beta_{\text{init}}$.

**Step** $k$. ($k \geq 1$) Find the smallest integer $i_k > 0$ such that with $\overline{\beta} = \eta^{-i_k}\beta_{k-1}$:

$\mathcal{L}(\overline{v}_k^{\overline{\beta}}) \leq \mathcal{L}(v_k) + [\nabla_{\boldsymbol{v}}\beta(\boldsymbol{v}_k)]^T[\overline{v}_k^{\overline{\beta}} - v_k] + \overline{\beta}\|v_k - \overline{v}_k^{\overline{\beta}}\|_2^2$ ▷ $\overline{v}_k^{\overline{\beta}}$ is a test parameter.

Set $\beta_k = \eta^{-i_k}\beta_{k-1}$ and compute: $\nabla_{\boldsymbol{v}}\mathcal{L}(\boldsymbol{v}_k)$,

$t_k = \frac{1+\sqrt{1+4t_k^2}}{2}$,

$\boldsymbol{v}_{k+1} = \boldsymbol{\gamma}_k + \frac{t_k - 1}{t_{k+1}}(\boldsymbol{\gamma}_k - \boldsymbol{\gamma}_{k-1})$.

**if** DistSupp($\boldsymbol{\gamma}_k$) $\neq$ Supp($\boldsymbol{\gamma}_k$) **then**

    UpdateSupp($\boldsymbol{\gamma}_k$)

In principle, this procedure should work when RoSE's PSF model matches the physical imaging system. However, by introducing such an adaptive step, the program in Alg. 1 is no longer convex and thus may not converge in a given iteration budget, especially in the face of model mismatch. To avoid possible instabilities, we can simply allocate half of our iteration budget to adjusting the support of the parameters and run the remaining iterations without updating the support, thereby making sure the program is convex and convergence is guaranteed. In the main paper and this SI, we refer to the (older) version with support update during all iterations (3) as **non-optimized RoSE** and the aforementioned hybrid version simply as **RoSE**. For a comparison of the two algorithms on experimental 3D SMLM data of microtubules, please refer to Fig. S22.

### 5. Supplementary Note 5: Background estimation

For the synthetic and experimental microtubule SMLM datasets (Figs. 2, 3), we used an iterative estimation algorithm based on the Wavelet transform with a wavelet level of 6 and the db6 basis family (10). For the TAB datasets (Figs. 4, 5, S25-S27), we averaged (spatially and temporally) the detected photons across all frames in a $20 \times 20$ pixel$^2$ region near the fibril but away from any Nile red blinking events. Assuming spatial uniformity, we used this average photon flux as our estimate of the background for localizing SMs (RoSE) and for computing WIF.

For the 3D microtubule dataset (Fig. 3), we first localize the Gaussian-like lobes of the DH-PSF images by using ThunderSTORM (2D Gaussian fitting with an initial $\sigma$ of 256 nm and a fitting radius of 640 nm). Next, we remove the detected signal photons from each SM by setting all pixels to NaN (not a number) within a square window (5-pixel side length) centered at each localization. Next, we fit each row and column of the image independently to a 1D Gaussian function, ignoring the NaN entries from the previous step, to obtain a non-smooth background estimate. Finally, we smooth the background using Wavelet filtering with a wavelet level of 4 and the db6 basis family.

Finally, in the special cases of images of isolated molecules (Fig. S22a-c) or beads (Fig. S21), we average nearby pixels (chosen by hand corresponding to background) in order to estimate the photon flux and background.

### 6. Supplementary Note 6: Localization confidence of an isolated molecule

We assess the performance of WIF by analyzing images of fluorescent molecules, generated using a vectorial image formation model (11), having various hidden physical parameters such as defocus and rotational mobility. As a baseline, we fix the PSF model in our confidence analysis to that of an isotropic molecule with zero defocus. To determine our confidence metric's robustness to shot noise, we use RoSE (3) to localize an isotropic emitter from 200 noisy, independent realizations of its image for a wide range of detected photons. Computing WIF for these localizations, we observe that the confidences are mostly close to 1 for all photon counts, taking values in [0.95, 1] (Fig. S4). There is a slight reduction in estimated confidences for large photon counts, most likely due to the first-order approximation in our PSF model (Supplementary Note 3, section D, Fig. S3). Furthermore, we conduct similar analysis for WIF3D using the DH-PSF (12). To this end, we use RoSE to analyze 300 noisy images of an isotropic emitter whose $z$-position is randomly chosen within $[-400, 400]$, which is fixed for all SNRs (Fig. S5). We observe that for all considered brightnesses, the median of confidences attain values above 0.8. In addition, as the brighntess increases, the standard deviations of the confidences decrease, which is expected. We note that the discrepancies between WIF2D and WIF3D may be due to our formulation of WIF3D.

Next, we quantify how hidden variables that are not accounted for within the model affect the confidences. For a dim molecule (800 photons and 20 background photons/pixel) at modest defocus values ($z \in [0, 200 \text{ nm}]$), we observe that the confidences mostly remain above 0.9 (Fig. S6a,b). As defocus increases beyond 200 nm, approximately 50% of localizations exhibit confidence lower than 0.9. In particular, for $z = 300$ nm, the median confidence decreases to 0.62, a reduction of approximately 40% from $z = 0$ (Fig. S6a). Our confidence metric is remarkably more sensitive to defocus compared to estimates of normalized PSF width (w.r.t. the PSF width at focus), which fluctuate mostly within 10% of their nominal values. For $z = 300$ nm, the median width reduces somewhat counter-intuitively by 13% from its nominal value (Fig. S6a), most likely because of the low SNR.

To explore how shot noise affects WIF and width estimates, we consider a bright molecule (2000 photons) (Fig. S6d). Interestingly, as soon as the defocus increases beyond 140 nm, the confidences sharply drop below 0.9 such that at $z = 200$ nm the median confidence approaches 0.3. In contrast, normalized width estimates remain mostly within 5% of their nominal values with their medians consistently close to 1 (Fig. S6c). Therefore, WIF even detects subtle defocus-induced model mismatches for brighter molecules with sufficient SNRs.

Next, we study how well WIF can quantify dipole-induced imaging errors, further exacerbated by defocus. We consider a molecule inclined at 45° with respect to the optical axis and with various degrees of rotational motion: effectively unconstrained or isotropic (uniform rotation within a cone of half angle $\alpha = 90°$), moderate confinement ($\alpha = 30°$), and strong constraint ($\alpha = 15°$) (Fig. S7c). For a photon count of 1000, notably, we observe consistent decreases in median confidences (below 0.85) for both $\alpha = 30°$ and $\alpha = 15°$ across all $z$, while for the isotropic molecule, the median confidence drops below 0.9 only for $z$ greater than 160 nm. In addition, confidences for $\alpha = 15°$ are smaller than those of $\alpha = 30°$, which shows our confidence metric's consistency, trending smaller as the degree of mismatch increases (Fig. S7a). On the other hand, normalized width estimates are practically indistinguishable for all $\alpha$ and $z$ values (Fig. S7b).

We next consider a brighter molecule (2000 photons) (Fig. S8c), and observe that confidences for both $\alpha = 30°$ and $\alpha = 15°$ significantly decrease below 0.5 for almost all $z$ positions (Fig. S8a). Surprisingly, the normalized width estimates for $\alpha = 30°$ and $\alpha = 15°$ converge to their nominal (in focus) value as $z$ approaches 200 nm (Fig. S8b).

Lastly, we validate WIF in 3D using the Double-Helix 3D PSF (DH-PSF) (12) and quantify 3D confidences in the presence of refractive index mismatch between the imaging medium and the sample (Table S2, Fig. S9). Concretely, we compute the DH-PSF model assuming a perfect match between the refractive indices of the medium and the sample (refractive index=1.51) and use this PSF model for 3D localizations using RoSE as well as for computing the WIFs or confidences. Next, we generate DH-PSF images with the sample refractive indices chosen within the range $[1.3, 1.51]$. Specifically, for each elected sample refractive index, we simulate 300 isolated images of a molecule (brightness 4000 photons) with its $z$-position randomly chosen within $[-400, 400]$ nm. We set the background to 20 photons per pixel and it is assumed to be known *a priori* in computing WIF.

When the refractive index of the medium is matched to that of the sample, we observe high confidences for RoSE's localizations (median 0.91, Fig. S9). As the sample refractive index decreases to 1.3, the median of the confidences also drops to 0.64, signaling inaccurate localizations. Indeed, examining the axial errors between the estimated $z$-positions and the ground-truth ones, we see a high correlation between the axial localization errors and the corresponding confidences (a Pearson correlation of 0.97).

### 7. Supplementary Note 7: Effect of SNR on the performance of WIF in detecting position and brightness inaccuracies

To study the limitations of WIF versus SNR, we first generate images of a SM at various SNRs (each 200 realizations) and localize them using RoSE. Next, we add an offset position error of 11.7 nm to the x position of localized molecules and feed them to the WIF algorithm. Intuitively, we expect that below certain SNR, the likelihood landscape becomes uninformative and thus causes the "regularized transport" to prefer sparser solutions whose transport trajectories are mostly aligned with the position of the original source. In particular, position errors that are comparable to the achievable localization precision, especially at low SNRs, cannot be detected by WIF (Fig. S10). In addition, as the SNR increases, WIFs converge to small asymptotic values slightly below 0 (Fig. S10). Note that 0 is expected as approximately half of the transport trajectories converge toward the unperturbed estimate.

We next vary the level of the offset error according to the achievable precision at each SNR. We find that when the offset (position) error is beyond three times the achievable localization precision, WIF is able to detect inaccurate localizations with a classification/detection accuracy greater than 80% using a WIF threshold of 0.5. The achievable localization precision is computed according to a formula in Ref. (13). In particular, the best lateral localization precision can be approximated as the square root of

$$\Delta x^2 = \frac{\sigma^2 + a^2/12}{s}\left(1 + 4\tau + \sqrt{\frac{2\tau}{1+4\tau}}\right), \text{ where} \qquad [41]$$
$$\tau = \frac{2\pi b(\sigma^2 + a^2/12)}{sa^2},$$

$a$ is the camera pixel size in object space, $s$ is the brightness of a molecule, $b$ is the background photons per pixel, and $\sigma$ is the width of the Gaussian spot. A similar expression for the best precision of brightness, i.e., $s$, can be derived (13):

$$\Delta s^2 = s\left(1 + 4\tau + \sqrt{\frac{\tau}{14(1+2\tau)}}\right). \qquad [42]$$

Next, we study the limitations of WIF versus SNR to quantify errors in estimating the expected brightness of a molecule. Such a scenario can happen due to various aberrations in the PSF. We consider two types of aberrations: defocus and astigmatism. To model defocus, we generated 200 images of a molecule located at (0,0,180 nm) with various brightnesses and a background of 20 photons per pixel (Fig. S11d). We used RoSE with a PSF computed at focus to localize molecules. As the brightness increases, the error in estimating the brightness also increases as the greater number of photons diffuse around the molecule. The results indicate WIF values significantly decrease below 0.5 as soon as the brightness error increases beyond 477 photons. Note that for high SNR, e.g., brightness of 7000 photons, WIF approaches -1. This is expected because all of the transport trajectories tend to diverge from the original localization, which is at the center of grid (Fig. S11c).

For astigmatism aberration, we generated images of molecule located at (0,0,50) nm with a small oblique astigmatism (Fig. S11h). We observe overall similar behavior for WIF as in the case of defocus aberration. However, there are some differences in the behaviour of WIF. First, for approximately same level of brightness error (e.g., 250 and 220 photons), WIFs are smaller for the astigmatism. This behaviour could be explained by observing that with defocus the effective SNR of the image is lower compared to images aberrated by astigmatism (Fig. S11d,h). Therefore, the transport trajectories are more informative in the case of astigmatism and thus produce more accurate WIFs. On the other hand, when a molecule is defocused and many of its photons have diffused away from its true location, its image essentially looks identical to that of a dim molecule. Unfortunately, these diffused photons are comparable to the background fluctuations surrounding the molecule. Therefore, for this particular scenario, a higher brightness, or SNR, is required for WIF to reliably detect these errors. Note that in the limit of high SNR, WIF approaches zero for astigmatism. This effect is expected as roughly half of the transport trajectories diverge from the original localization (Fig. S11g).

Interestingly, the regularized transport trajectories can serve as useful diagnostic tools to decipher the cause of low WIFs. For example, in the case of defocus aberration, all trajectories diverge (Fig. S11c), while for oblique astigmatism only those perturbed sources that lie parallel to the elongated PSF axis diverge (Fig. S11g).

### 8. Supplementary Note 8: Quantifying localization accuracy via $\mathrm{WIF}_{\mathrm{avg}}$

We simulate SMLM datasets of stochastically-blinking molecules using an ideal imaging model, i.e, with no mismatch ( Table S2, Fig. S16a). For each frame we generate images with a certain blinking density (defined as the number of active molecules per $\mu\mathrm{m}^2$) by randomly selecting the positions of molecules in the field-of-view. As proposed in the main text, for a set of $N$ localizations returned by an algorithm with confidences $\{c_1, \ldots, c_N\}$, we use $\mathrm{WIF}_{\mathrm{avg}} \triangleq \frac{1}{N}\sum_{i=1}^{N} c_i$ to quantity the collective accuracy of the said algorithm. We can gain insight into $\mathrm{WIF}_{\mathrm{avg}}$ by examining its correspondence to the well-known Jaccard index (JAC), which determines the credibility of a localization based on its distance to the ground-truth SM. In particular, we may define $\mathrm{JAC} = \mathrm{TP}/(\mathrm{TP}+\mathrm{FN}+\mathrm{FP})$, where TP, FN, and FP denote number of true positives, false negatives, and false positives, respectively. An undetected molecule, that is, a false negative, would increase the denominator of JAC, thereby reducing its value. We posit that this same undetected molecule adversely affects the confidence of a nearby localized molecule, thereby reducing $\mathrm{WIF}_{\mathrm{avg}}$. This intuitive connection between JAC and $\mathrm{WIF}_{\mathrm{avg}}$ suggests that the average confidence may serve as a good surrogate for localization accuracy.

Using $\mathrm{WIF}_{\mathrm{avg}}$, we quantify the performance of three algorithms, RoSE (3), FALCON (8), and ThunderSTORM (TS) (6), for localizing emitters at various blinking densities (Fig. S16a,b). Examining the localizations returned by the algorithms, we calculate the Jaccard index using ground-truth information from an oracle and $\mathrm{WIF}_{\mathrm{avg}}$ using only the simulated images of SM blinking. For all RoSE, FALCON, and TS, we observe excellent agreement between $\mathrm{WIF}_{\mathrm{avg}}$ and Jaccard index for densities as high as 5 mol./$\mu\mathrm{m}^2$. For higher densities, $\mathrm{WIF}_{\mathrm{avg}}$ monotonically decreases at a rate differing from that of Jaccard index. For instance, at high densities JAC for TS saturates to 0.1, whereas $\mathrm{WIF}_{\mathrm{avg}}$ further decreases due to high FN and low TP (Fig. S16c), thus penalizing its poor performance.

A natural application of our confidence metric is to remove localizations with poor accuracy. We filter localizations with confidence smaller than 0.5, corresponding to half of the perturbed photons “returning” toward a particular localization, and calculate the resulting $\mathrm{precision} = \mathrm{TP}/(\mathrm{TP}+\mathrm{FP})$ and $\mathrm{recall} = \mathrm{TP}/(\mathrm{TP}+\mathrm{FN})$. If the filtered localizations truly represent false positives, we expect to see an increase in precision and a relatively unchanged recall after filtering. Our results show a precision enhancement as high as 180% for TS and a desirable increase of 23% for RoSE (density= 9 mol./$\mu\mathrm{m}^2$) (Fig. S16d). Remarkably, these improvements come with a negligible loss in recall (13% in the worst case) across all densities for both algorithms (Fig. S16e).

We applied similar analyses for the 3D DH-PSF. We used RoSE to localize molecules at various blinking densities (Fig. S17). Here, we define the density as the number of molecules per volume (5 $\mu\mathrm{m}^3$, $2.5 \times 2.5 \times 0.8$ $\mu$m). Upon removing localizations with $\mathrm{WIF} < 0.5$, we note that the performance loss in Recall in Fig. S17 is higher compared to that of 2D case (Fig. S16). This loss may be due to limitations imposed by the DH-PSF, a small value of $\nu$ in computing WIFs (Supplementary Note 3C), or a large distance tolerance for assigning false positives (here it is set to 44 nm).

### 9. Supplementary Note 9: Pupil fitting

Recall that, in principle, we can use Fourier optics to model the microscope PSF $q$ as the Fourier transform of a generalized pupil function (14). In particular, for a specific axial position $z$, we can compute the corresponding PSF $q(z)$ as:

$$q(z) = |\mathcal{F}\{P(z)\}|^2, \qquad [43]$$

where $\mathcal{F}$ represents the Fourier transform, $P(z)$ represents the microscope pupil evaluated at $z$, and $|\cdot|$ denotes the element-wise absolute value operator. We may further simplify Eq. (43) by assuming a global pupil $P_0$, which is modulated by a defocus term: $P(z) = P_0 \cdot \text{defocus}(z)$.

For a given set of $K$ images of a bead sampled at specific axial positions, we may formulate an optimization problem in which we aim to fit $P_0$. Here, we consider the following optimization

$$\hat{P}_0 = \operatorname*{arg\,min}_{P_0 \in \mathbb{R}^{N\times N}} \sum_{i=1}^{K} \mathcal{L}(g_i; P_0), \qquad [44]$$

where $N$ denotes the size of the microscope pupil (in pixels), $g_i \in \mathbb{R}^{m\times m}$ represents the $i^{\text{th}}$ image of a bead at an axial position $z_i$, and $\mathcal{L}$ is the Poisson negative log likelihood. We used an automatic differentiation package (autograd Version 1.3 (15)) in Python to estimate $P_0$. The resulting PSF model is computed by plugging $\hat{P}_0$ into Eq. (43).

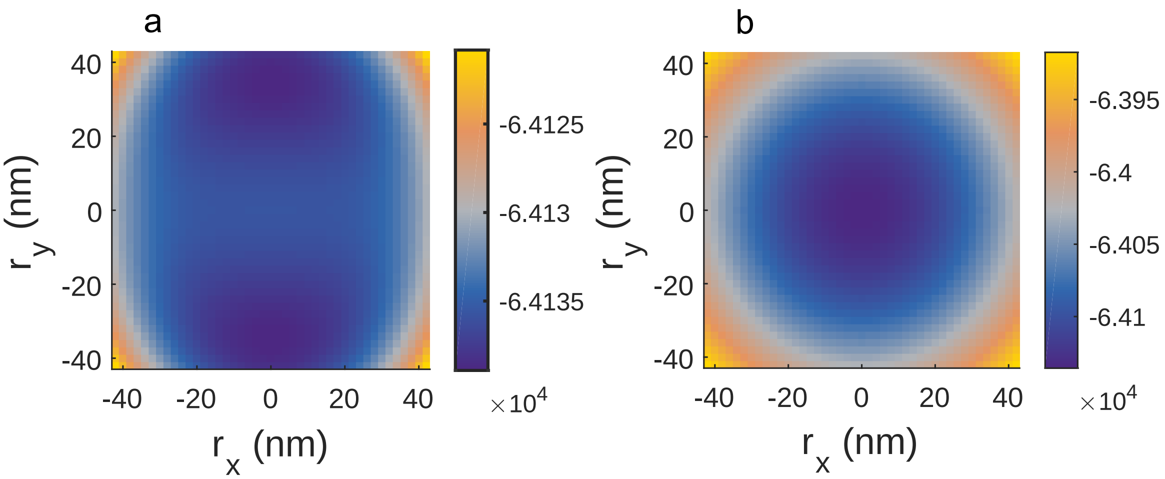

**Fig. S1.** Landscape of expected, negative Poisson log likelihood in localizing two closely-spaced molecules. (s) Expected, negative Poisson log likelihood for a model parameterized by two equally bright molecules, which are located at $(r_x, r_y)$ and $(-r_x, -r_y)$. (b) Similar to (a), but for a model parameterized by one molecule, which is located at $(r_x, r_y)$. Ground-truth molecules are located at $(0, -35)$ and $(0, 35)$ nm with equal brightnesses of $2000$ photons. Noisy images were generated according to a symmetric Gaussian PSF with $\sigma = 0.21\lambda/\text{NA} = 100$ nm. Background was set to $20$ photons per pixel.

 

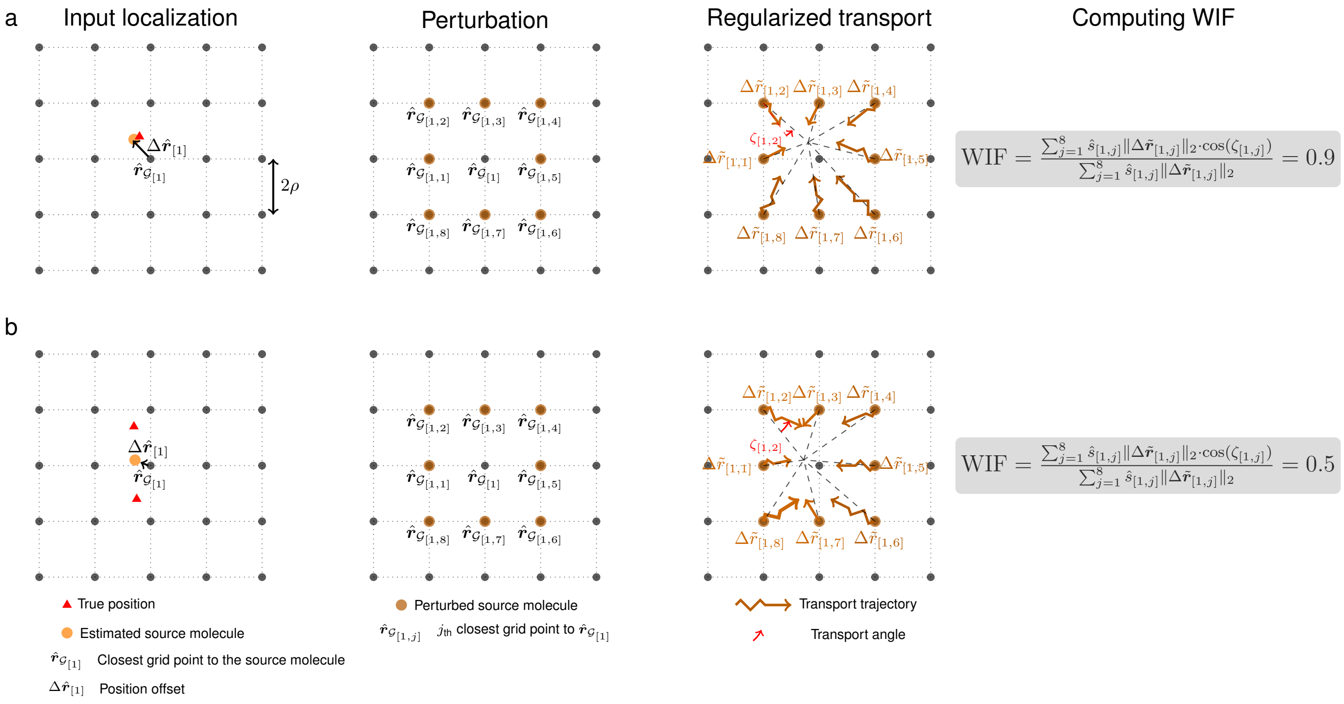

**Fig. S2.** Computing Wasserstein-induced flux (WIF). (a) (left) Input localization (orange circle) located at $\hat{\boldsymbol{r}}_{[1]}$ is mapped to its closest grid point $\hat{\boldsymbol{r}}_{\mathcal{G}_{[1]}}$ and a position offset vector $\Delta\hat{\boldsymbol{r}}_{[1]}$, i.e., $\hat{\boldsymbol{r}}_{[1]} = \hat{\boldsymbol{r}}_{\mathcal{G}_{[1]}} + \Delta\hat{\boldsymbol{r}}_{[1]}$ . Note that $2\rho$ indicates the distance between any two points on the grid. (middle left) A perturbation redistributes a molecule's photons to its 8 closest grid points $\{\hat{\boldsymbol{r}}_{\mathcal{G}_{[1,1]}}, \ldots, \hat{\boldsymbol{r}}_{\mathcal{G}_{[1,8]}}\}$. (middle right) Solving Eq. (36) amounts to finding a set of transport trajectories or displacements denoted by $\{\Delta\tilde{\boldsymbol{r}}_{[1,1]}, \ldots, \Delta\tilde{\boldsymbol{r}}_{[1,8]}\}$. The transport angle $\zeta_{[1,j]}$ is defined as the angle between the displacement $\Delta\tilde{\boldsymbol{r}}_{[1,j]}$ and $(\hat{\boldsymbol{r}}_{[1]} - \hat{\boldsymbol{r}}_{\mathcal{G}_{[1,j]}})$. (right) WIF is computed according to the recipe described in Eq. (32). Note that since the estimated localization (orange circle) is close to the true position (red triangle), we expect WIF to be close to 1, indicating a high degree of confidence. (b) Similar to (a), but for an inaccurate input localization (orange circle). Note that the computed WIF is much smaller than 1, thereby signaling a high degree of uncertainty.

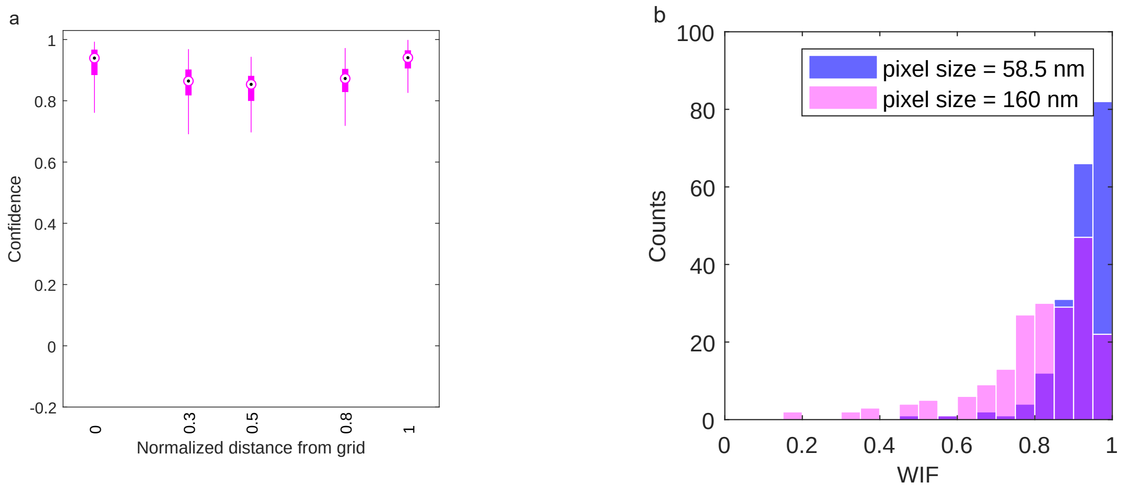

**Fig. S3.** Effect of PSF approximation on WIF as a function of the molecule's position relative to the computational grid and camera pixel size. (a) Box-plot of confidences/WIFs of a molecule at various distances from a grid point located at (0,0). Black circles represent the medians of confidence distributions, while filled boxes (magenta) indicate the range of $25^{\text{th}}$ to $75^{\text{th}}$ percentiles. Here, the normalized distance of 1 means that the location of the molecule is $(58.5/2, 58.5/2)$ nm, which is a neighbouring grid point. Note that $58.5$ nm is the camera pixel size or twice the grid distance. The brightness of the molecule is set at 3000 photons with a background of 20 photons per pixel and 200 realizations were used at each distance. (b) Effect of camera pixelation on WIF. For each camera pixel size, 58.5 nm and 160 nm, we simulated 200 independent images of a molecule located at the origin. For both cases, we used an expected brightness of 2000 photons. For pixel size of 58.5 nm, we used an average, uniform background of 20 photons per pixel, while for 160 nm pixel size, we used a uniform background of 54.7 photons per pixel, which ensures that the background level is appropriately scaled with the camera pixel size. To compute WIF, we used the same standard PSF model that we used to localize these molecules, and the grid distances for both pixel sizes were roughly the same (grid distances of 30 nm and 40 nm for 58.5 nm and 160 nm pixel sizes, respectively). We used a regularizer value of 0.1 for both cases. We note that the achievable localization precision (Ref. (16)) of $x$ and $y$ for both cases are virtually the same ($8.02$ nm versus $8.08$ nm), indicating that the difference in the WIF distributions is due to first-order approximation errors. Median WIFs: (58.5 nm pixel size) 0.94 and (160 nm pixel size) 0.84.

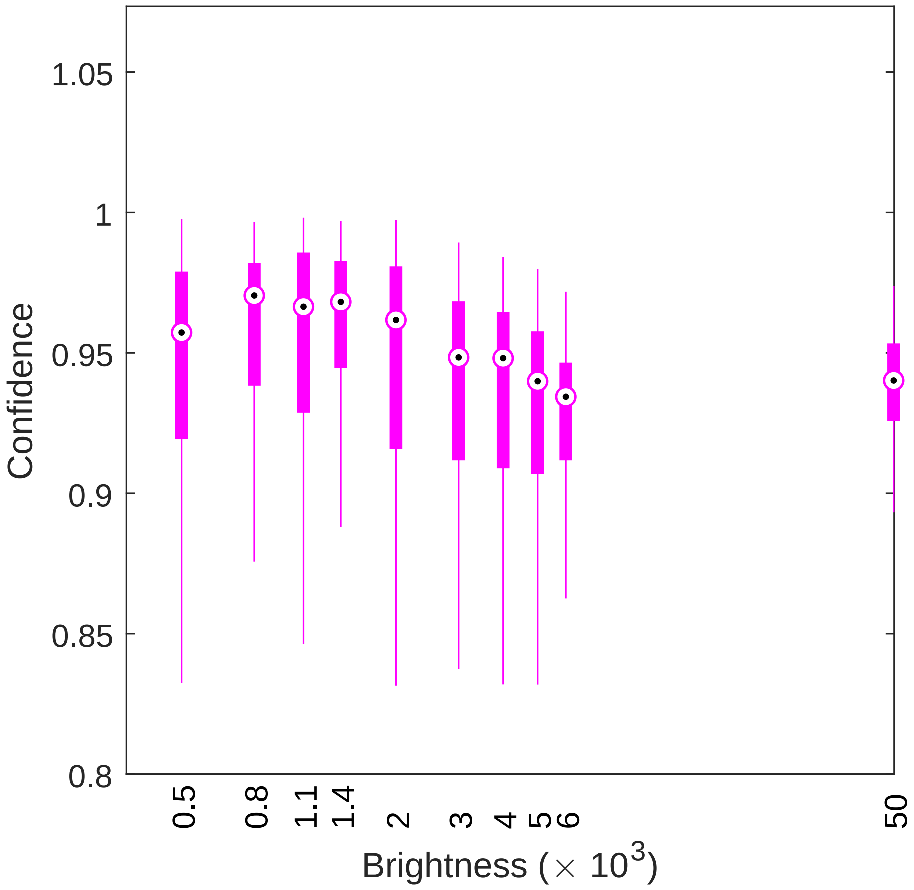

**Fig. S4.** 2D localization confidence of a molecule with various brightnesses. Black circles represent the medians of confidence distributions, while filled boxes (magenta) indicate the range of $25^{\text{th}}$ to $75^{\text{th}}$ percentiles. For each brightness, $200$ independent images of an isotropic molecule were localized using RoSE and then analyzed. Background was set to $20$ photons per pixel.

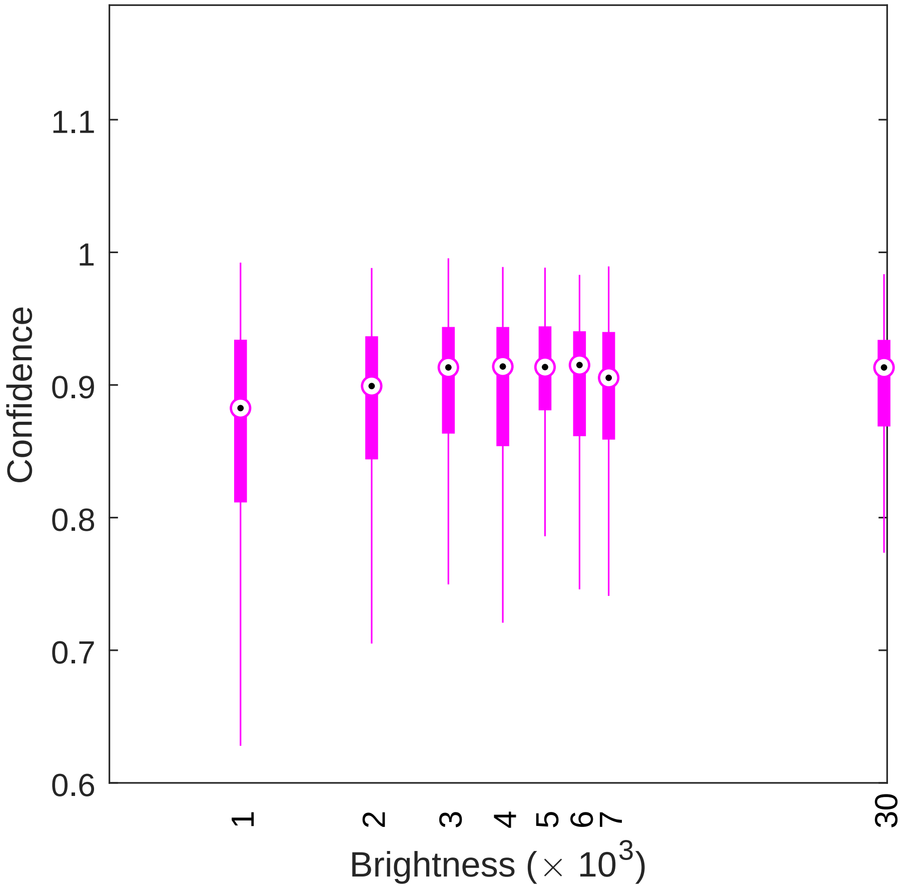

**Fig. S5.** 3D localization confidence of a molecule with various brightnesses. Black circles represent the medians of confidence distributions, while filled boxes (magenta) indicate the range of $25^{\text{th}}$ to $75^{\text{th}}$ percentiles. For each brightness, $300$ independent images of an isotropic molecule using the DH-PSF with a fixed but randomly-chosen z-position were localized using RoSE and then analyzed. Background was set to $20$ photons per pixel. The confidence analysis algorithm used an ideal model of the DH-PSF for an isotropic emitter.

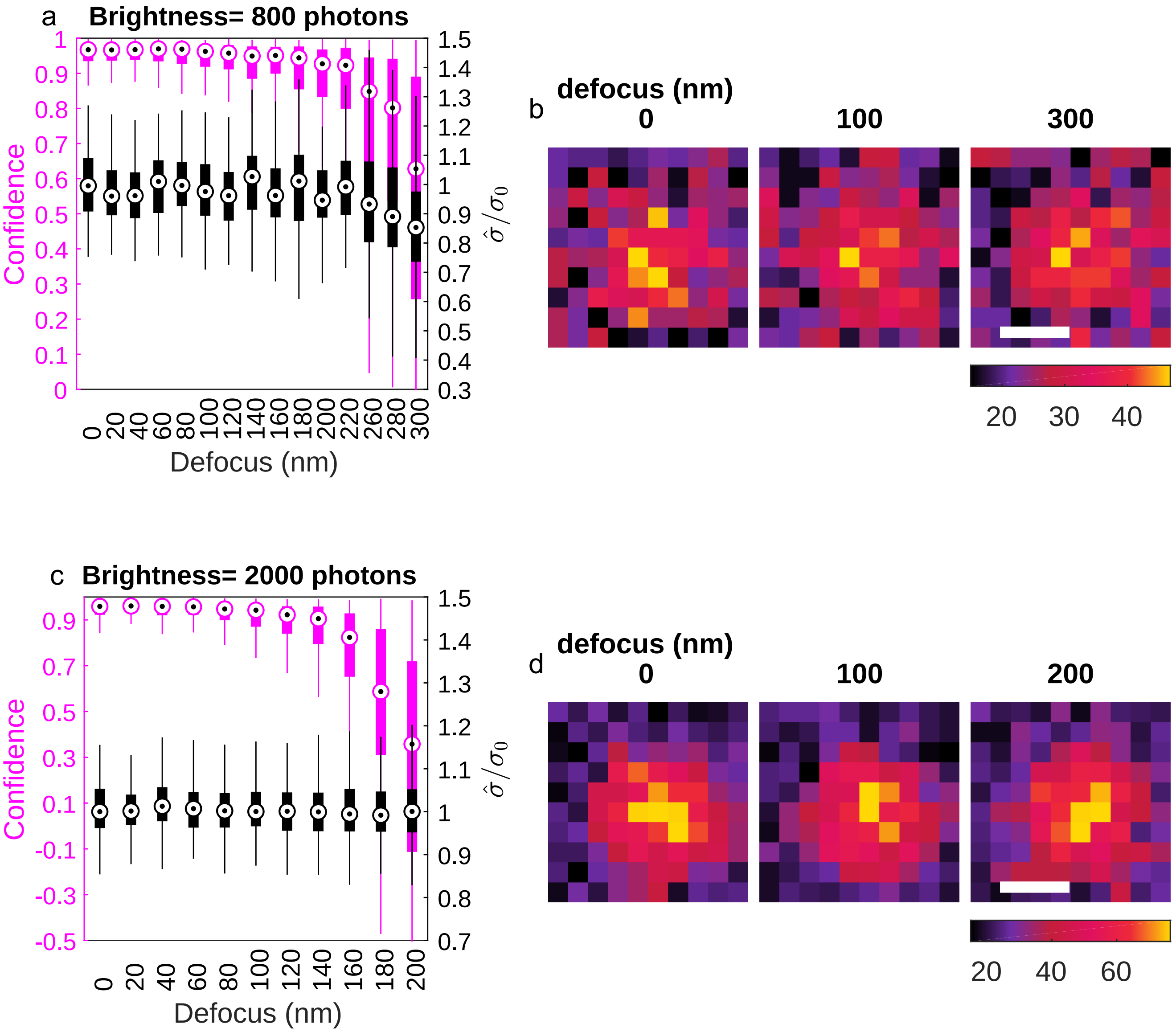

**Fig. S6.** 2D localization confidence of a molecule with various defocus mismatches. (a) Distributions of confidence (magenta) and normalized PSF width (black, $\hat{\sigma}/\sigma_0$). Black circles represent medians while filled boxes indicate the range of $25^{\text{th}}$ to $75^{\text{th}}$ percentiles. (b) Examples of images analyzed in (a) for a defocus of $0$ (left), $100$ nm (middle), and $200$ nm (right). (c) Similar to (a), but for a molecule with a brightness of $2000$ photons. (d) Similar to (b), but for images analyzed in (c). For each defocus value, $200$ independent images of an isotropic molecule with a brightness of $800$ photons were localized using RoSE and then analyzed. Background was set to $20$ photons per pixel. The confidence analysis algorithm used an ideal PSF model for an isotropic emitter with zero defocus. Colorbars: (b,d) photons$/58.5 \times 58.5$ nm$^2$. Scalebars: (b,d) $200$ nm.

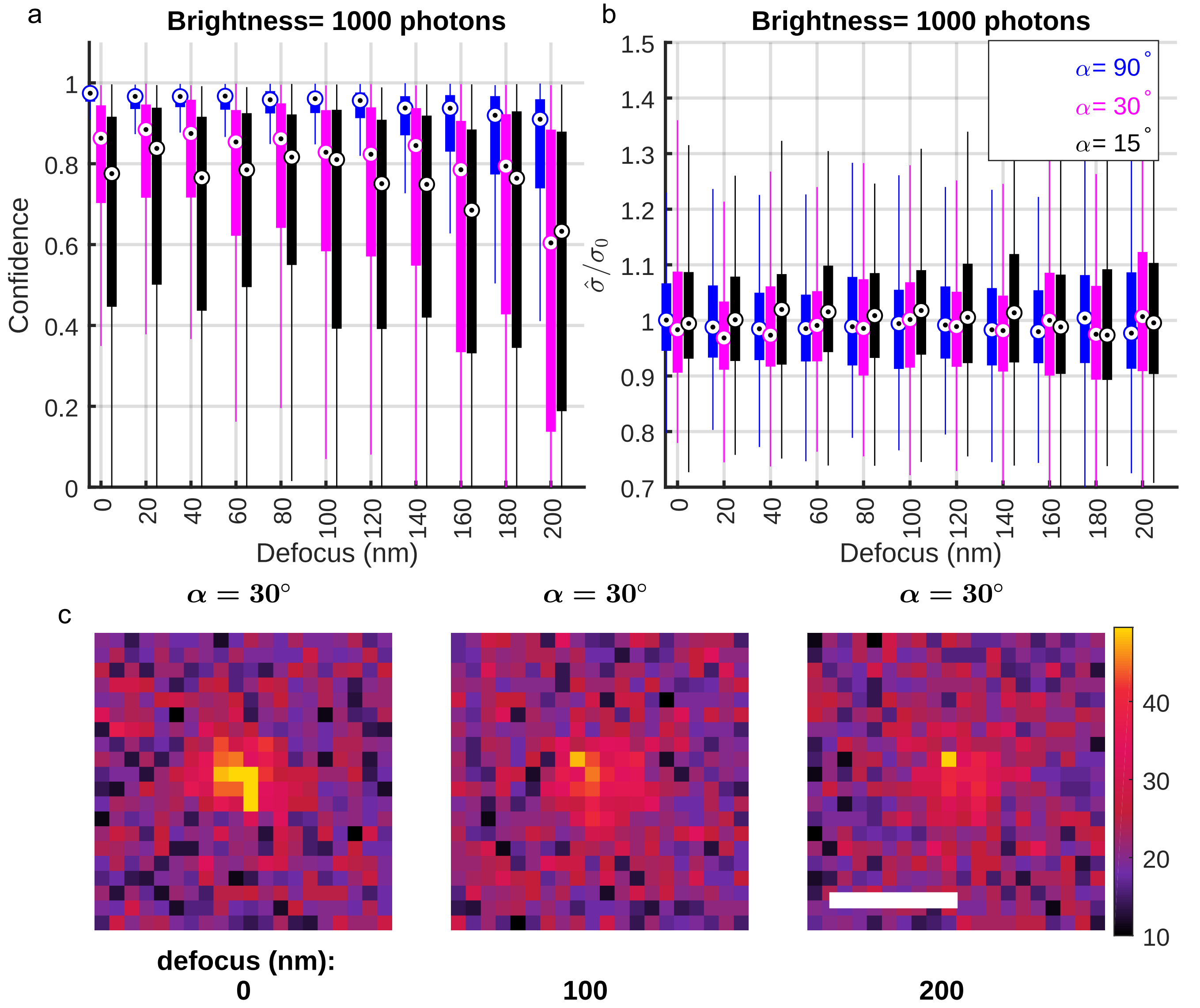

**Fig. S7.** 2D localization confidence of a molecule with various defocus and dipole-induced mismatches. (a) Distributions of confidence. Black circles represent medians, while the filled boxes (blue: uniform orientational diffusion within a cone of half-angle $\alpha = 90°$, magenta: $\alpha = 30°$, and black: $\alpha = 15°$ ) indicate the range of $25^{\text{th}}$ to $75^{\text{th}}$ percentiles. (b) Similar to (a), but for normalized PSF width estimates ($\hat{\sigma}/\sigma$). (c) Examples of images analyzed in (a,b) for $\alpha = 30°$ and a defocus of $0$ (left), $100$ nm (middle), and $200$ nm (right). For each defocus value, $200$ independent images of a dipole, each with a brightness of $1000$ photons and rotating uniformly with an polar angle of $45°$ and an azimuthal (in-plane) angle of $0$, were localized using RoSE and then analyzed. Colorbar: (c) photons$/58.5 \times 58.5$ nm$^2$. Scalebar: (c) $500$ nm.

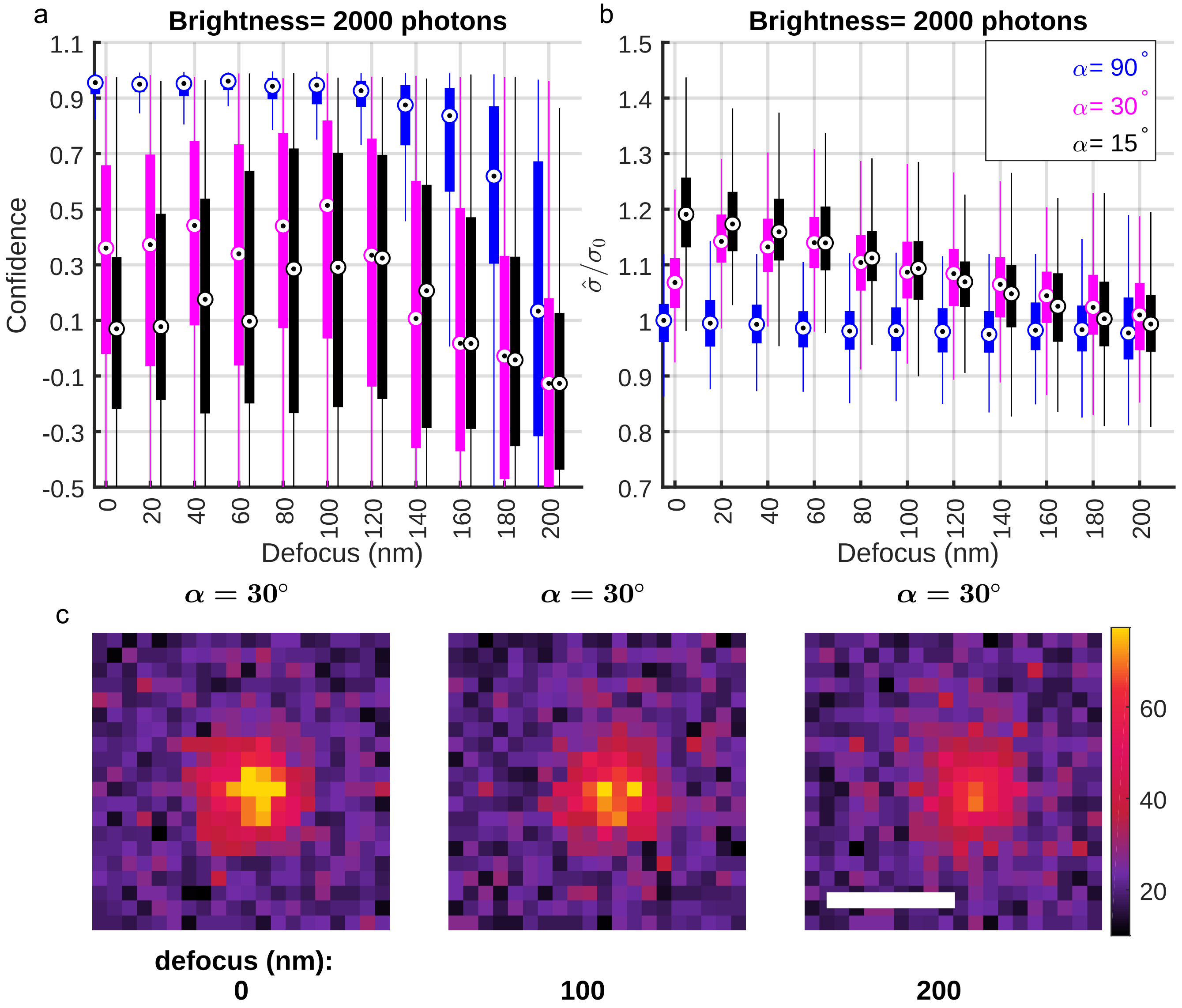

**Fig. S8.** 2D localization confidence of a bright molecule with various defocus and dipole-induced mismatches. (a) Distributions of confidence. Black circles represent medians, while the filled boxes (blue: uniform orientational diffusion within a cone of half-angle $\alpha = 90°$, magenta: $\alpha = 30°$, and black: $\alpha = 15°$ ) indicate the range of $25^{\text{th}}$ to $75^{\text{th}}$ percentiles. (b) Similar to (a), but for normalized PSF width estimates ($\hat{\sigma}/\sigma$). (c) Examples of images analyzed in (a,b) for $\alpha = 30°$ and a defocus of $0$ (left), $100$ nm (middle), and $200$ nm (right). For each defocus value, $200$ independent images of a dipole, each with a brightness of $2000$ photons and rotating uniformly with an polar angle of $45°$ and an azimuthal (in-plane) angle of $0$, were localized using RoSE and then analyzed. Colorbar: (c) photons$/58.5 \times 58.5$ nm$^2$. Scalebar: (c) $500$ nm.

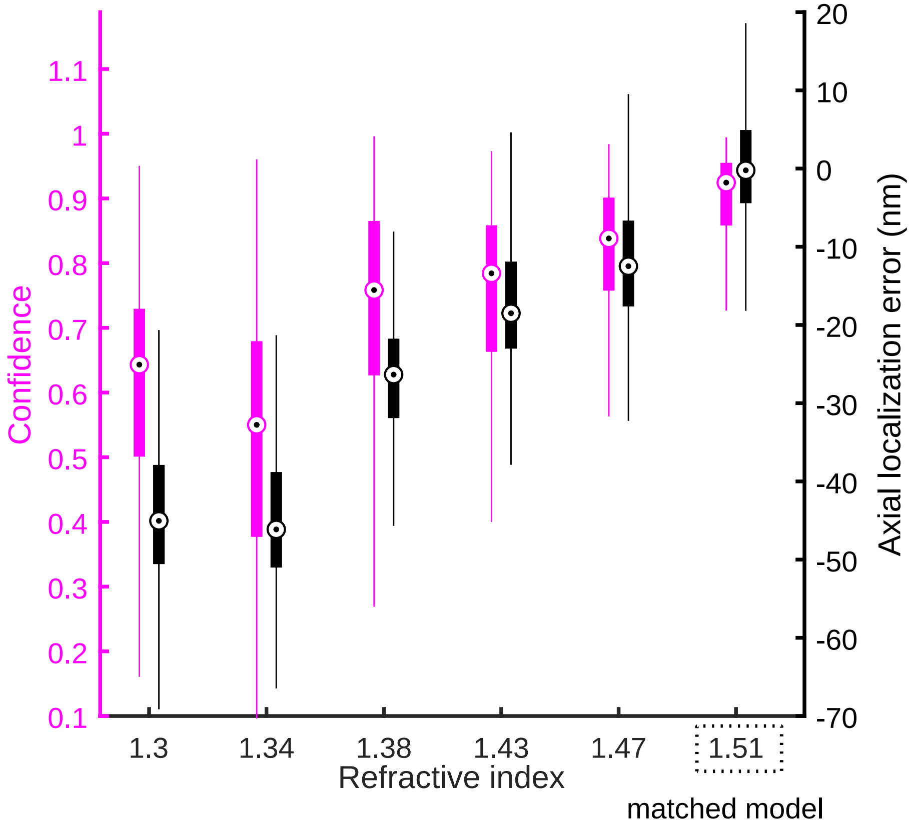

**Fig. S9.** 3D localization confidence and axial localization error of a molecule in a medium of mismatched refractive index. Black circles represent medians, while the filled boxes (black: axial localization error, magenta: 3D localization confidence) indicate the range of $25^{\text{th}}$ to $75^{\text{th}}$ percentiles. For each value of the sample's refractive index, $300$ independent images of an isotropic molecule with a brightness of $4000$ photons and a random z-position chosen within $[-400, 400]$ nm were localized using RoSE and then analyzed. Background was set to $20$ photons per pixel. The confidence analysis algorithm used an ideal model of the DH-PSF for an isotropic emitter within a matched refractive index.

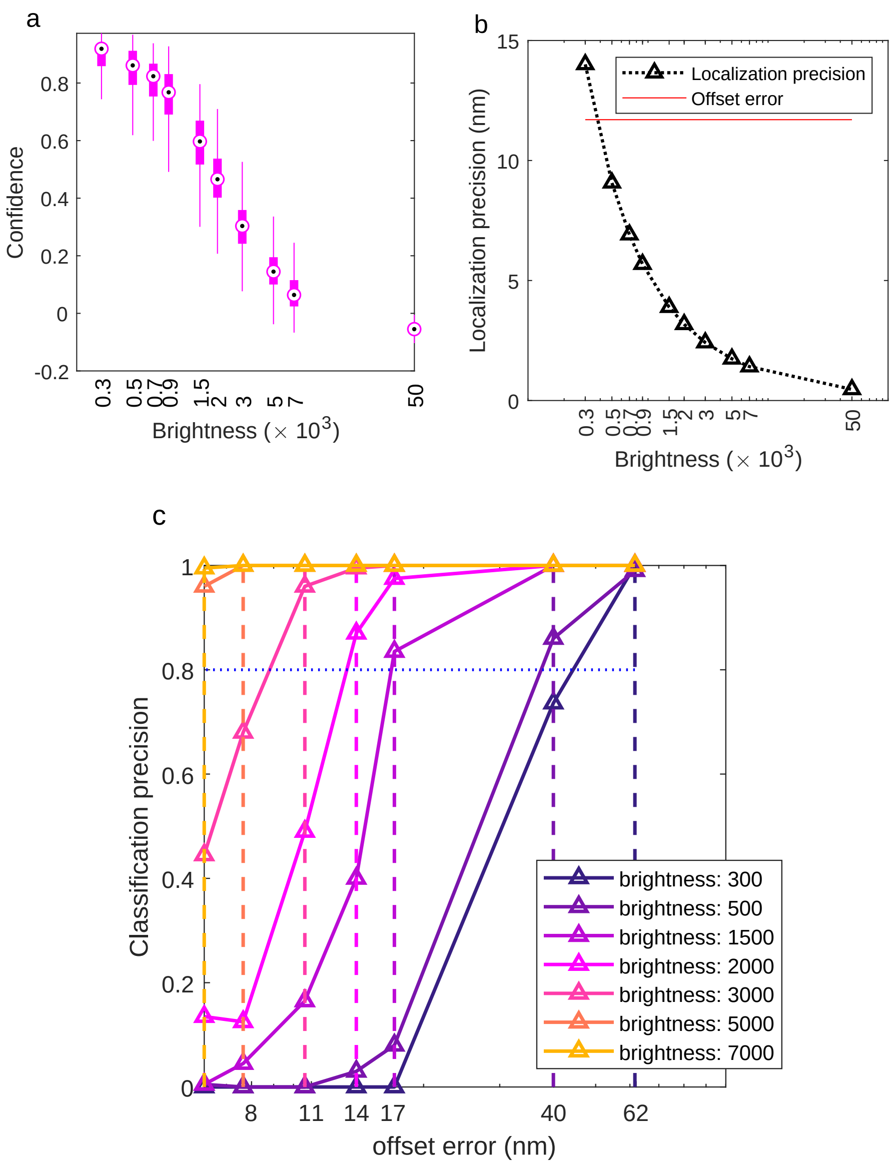

**Fig. S10.** Effect of SNR on the performance of WIF in quantifying position inaccuracy. (a) Box-plot of confidences/WIFs for localizations (200 at each SNR) with an offset error of $11.7$ nm for various brightnesses. Black circles represent medians, while the filled boxes indicate the range of $25^{\text{th}}$ to $75^{\text{th}}$ percentiles. (b) Achievable localization precisions at each SNR using Eq. (41) from Ref. (13). The red line indicates the value of the offset error (11.7 nm) added to each localization. (c) Precision of using a WIF threshold of 0.5 as an indicator of accurate localizations as a function of offset error. Precision = true positives/(true positives + false positives). All localizations have a non-zero position offset error; thus, localizations with a WIF score of less than 0.5 are assigned as true positives. The vertical dashed lines indicate three times the values of the achievable localization precision at each SNR. The dotted, horizontal line represents a (classification) precision of $0.8$ across all SNRs. Background for all SNRs was set to 20 photons per pixel; camera pixel size in object space was set to $58.5$ nm; and the width of the standard PSF was set to 95.5 nm.

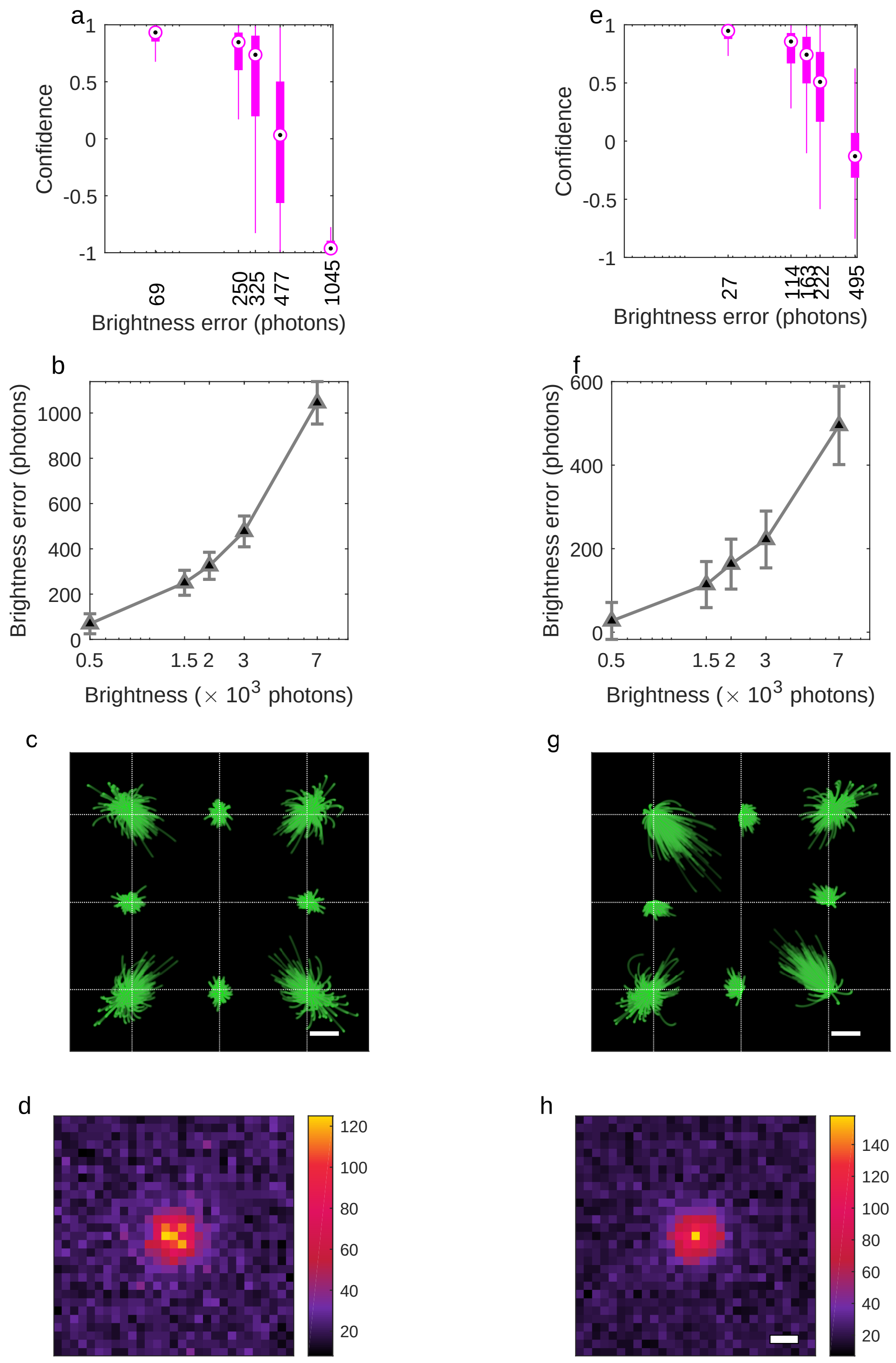

**Fig. S11.** Effect of SNR on performance of WIF in quantifying brightness errors due to aberrated PSFs. (a) Box-plot of confidences/WIFs for molecules (200 at each SNR) with defocus-aberrated PSFs that cause various brightness errors during localization. Note that the PSF model used to calculate WIF is obtained at focus. Black circles represent medians, while the filled boxes indicate the range of $25^{\mathrm{th}}$ to $75^{\mathrm{th}}$ percentiles. (b) Brightness errors in (a) plotted against their corresponding true brightnesses. The error bars represent the achievable brightness precision at each SNR. (c) Transport trajectories of all 200 realizations for defocus aberration in (a). (d) Representative images of a defocus aberration at a brightness of 3000 photons and 20 background photons per pixel. (e-h) Similar to (a-d), but obtained for oblique astigmatism aberration. Colorbar: photons$/58.5 \times 58.5$ nm$^2$. Scale bars: (c,g) 10 nm; (h) 200 nm.

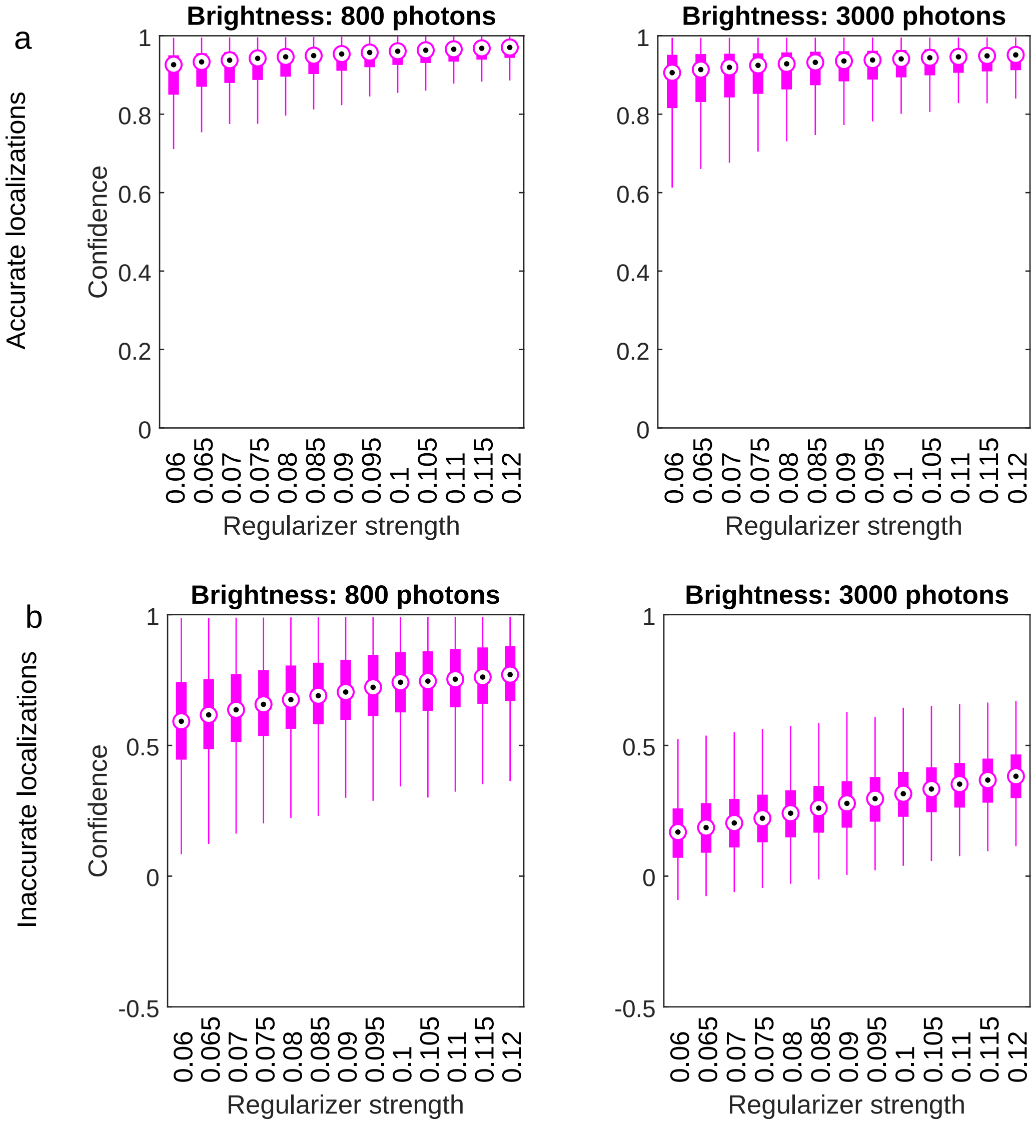

**Fig. S12.** Effect of regularizer strength on WIF for accurate and inaccurate localizations. (a) WIFs/confidences for accurate localizations (200 realizations at each regularizer strength) verus regularizer strength at brightness of (left) 800 and (right) 3000 photons and 20 background photons per pixel. Black circles represent medians, while the filled boxes indicate the range of $25^{\text{th}}$ to $75^{\text{th}}$ percentiles. (b) Similar to (a), but for inaccurate localizations obtained by adding an offset error of 11.7 nm to the measured localizations in (a).

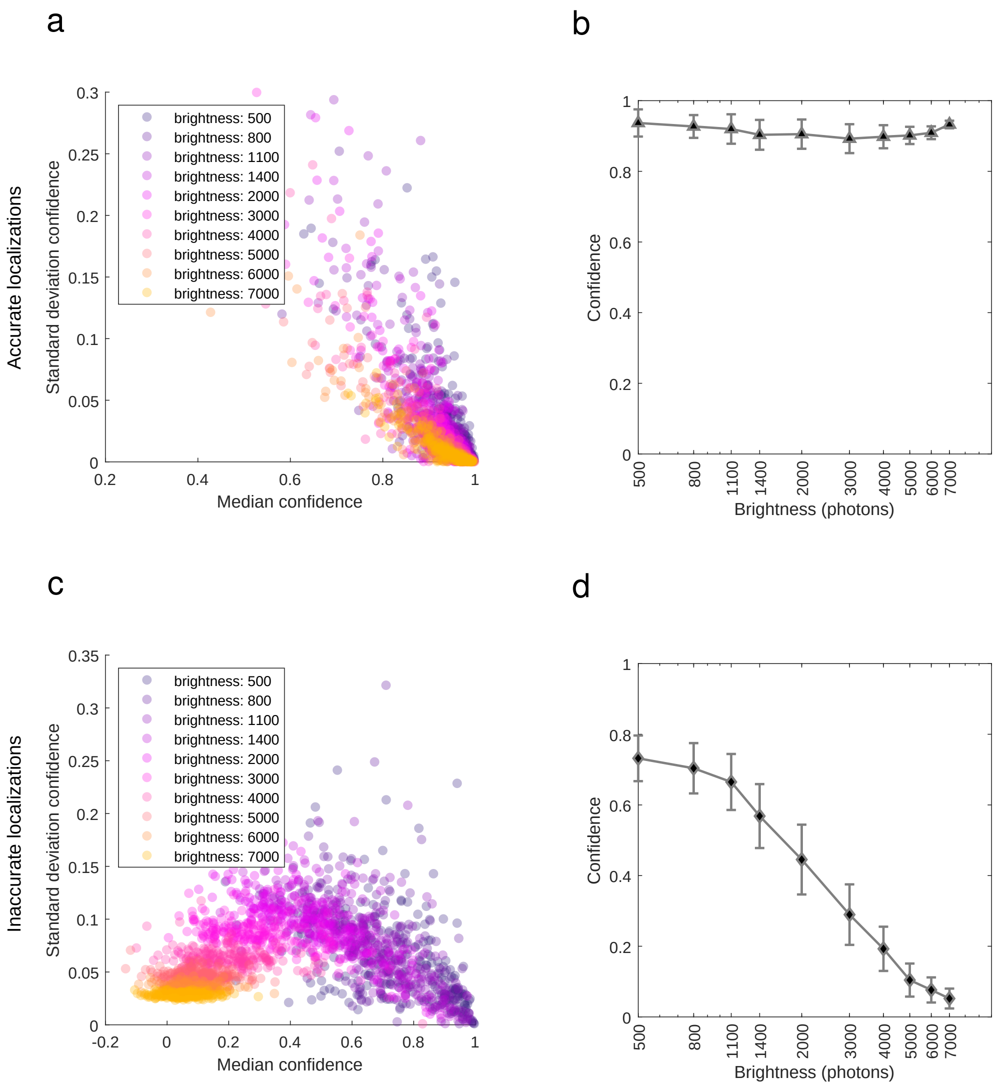

**Fig. S13.** Stability of WIF over a range of regularizer strengths for accurate and inaccurate localizations as a function of SNR. (a) Median vs. standard deviation of WIF for accurate localizations obtained at various brightnesses (200 realizations per each SNR). Standard deviations were obtained by calculating the median absolute deviations (MAD) of WIFs obtained for regularizer strengths of $[0.06, 0.065, 0.07, \ldots, 0.115, 0.12]$ (see Supplementary Note 3E). (b) Mean of WIF medians (black triangle) and $\pm$ mean of WIF standard deviations (error bars) of measured confidences for accurate localizations in (a). (c,d) Similar to (a,b), but for inaccurate localizations obtained by adding an offset error of 11.7 nm to the x positions of accurate localizations obtained in (a). For all SNRs background was set to 20 photons per pixel.

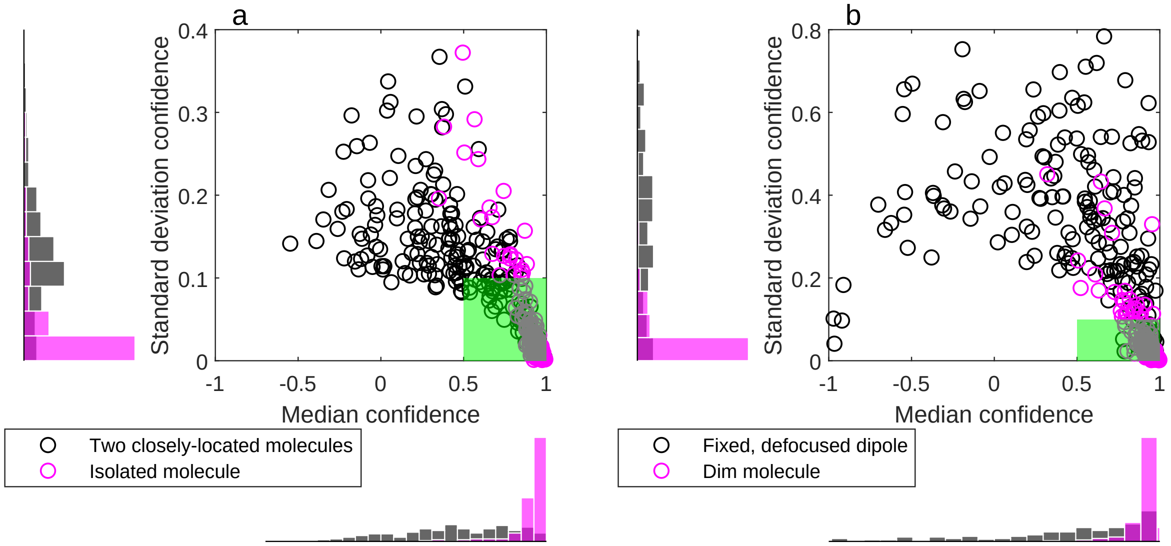

**Fig. S14.** Using median and standard deviation of WIFs calculated over a range of regularizer strengths significantly improves detection of localization inaccuracies. (a) Median vs. standard deviation of WIF for accurate localizations (200 realizations) of an isolated molecule (magenta) and inaccurate localizations of two-closely located molecules (black) (similar to Fig. 1c(i,ii)). The regularizer strengths used to calculate median and standard deviation WIFs are $[0.06, 0.065, 0.07, \ldots, 0.115, 0.12]$ (see Supplementary Note 3E). Brightness and background were set to 2000 photons per molecule and 20 photons per pixel, respectively. (b) Similar to (a), but for accurate localizations (200 realizations) of a dim molecule (magenta) and inaccurate localizations of a fixed, defocused dipole molecule (black) (similar to Fig. 1c(iii,iv)). Brightnesses were set to 2000 photons for dipole molecule and 800 photons for the dim molecule, and the background was set to 20 photons per pixel. The green, shaded box represents localizations that have a WIF median greater than $0.5$ and a WIF standard deviation less than $0.1$, which corresponds to the thresholds for classifying accurate versus inaccurate localizations mentioned in the main text.

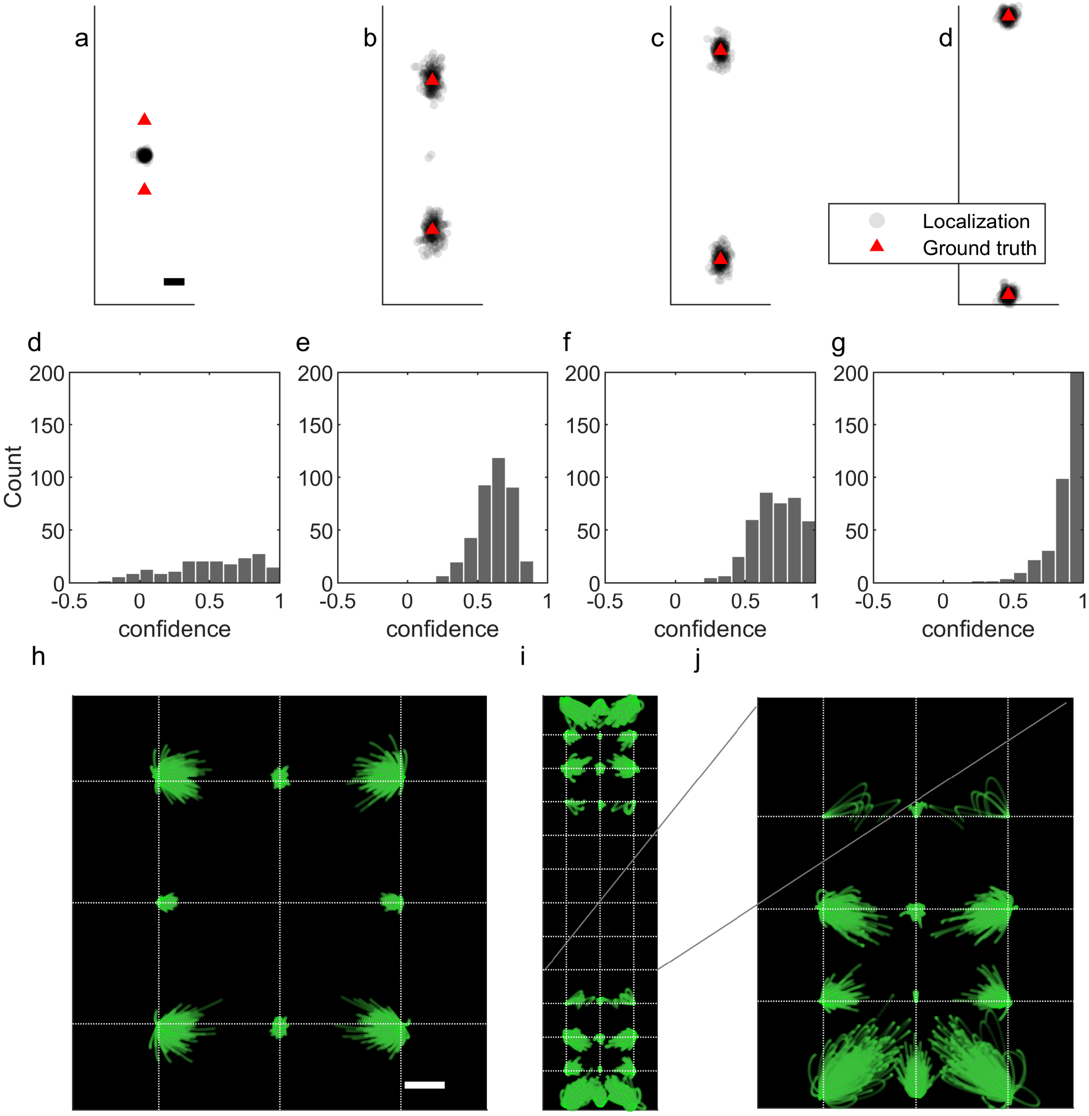

**Fig. S15.** WIF confidences for two closely-located molecules at various separation distances. Black circles indicate localizations obtained for 200 realizations of two molecules separated along y-axis by (a) 70 nm, (b) 150 nm, (c) 210 nm, and (d) 280 nm. The red triangles indicate the ground-truth position of the molecules. The brightness used in generating images is 2000 photons with 20 photons per pixel for background. (d,e,f,g) Corresponding WIFs for localizations in (a,b,c), and (d). (h) Transport trajectories for all 200 realizations in (a). (i) Similar to (h), but for localizations in (d). (j) Magnified version of the lower region in (i). Scalebars: (a) 20 nm, (h) 10 nm.

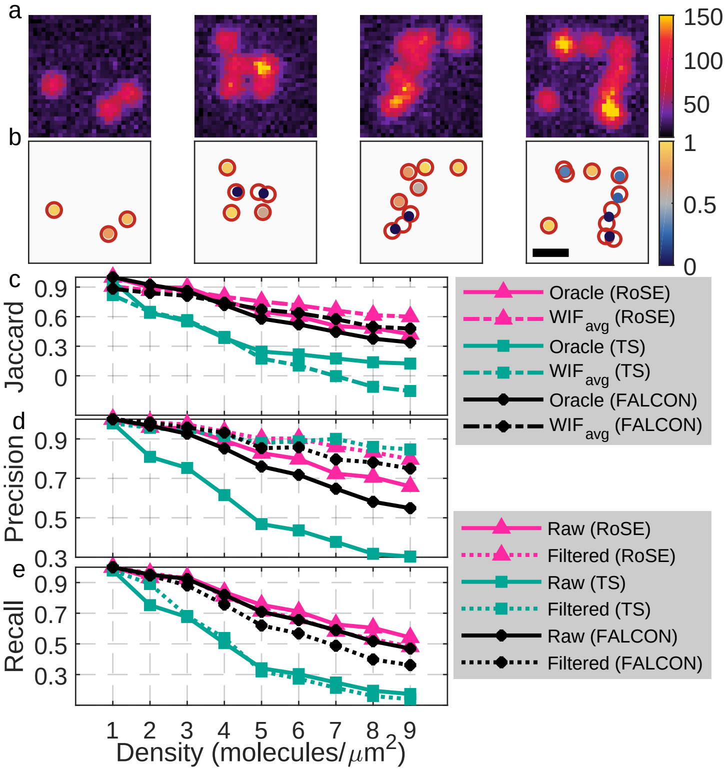

**Fig. S16.** Wasserstein-induced flux ($\mathrm{WIF_{avg}}$) quantifies localization accuracy without ground truth. (a) From left to right: images of molecules for blinking densities of 3, 5, 7, and 9 mol. per $\mu$m$^2$, respectively. (b) RoSE localizations (colored dots represent calculated confidence) corresponding to images in (a). Open red circles represent ground-truth positions. (c) Jaccard index (ratio of true positives to true positives, false positives, and false negatives) for RoSE (solid, red) and TS (solid, green) at various blinking densities. The dashed lines represent $\mathrm{WIF_{avg}}$ for RoSE (red) and TS (green). For each blinking density, 200 independent realizations were used. (d) Precision (ratio of true positives to true positives and false positives) for all localizations (solid) and localizations with confidence greater than $0.5$ (dotted) using RoSE (red) and TS (green). (e) Recall (ratio of true positives to true positives and false negatives) for all localizations (solid) and localizations with confidence greater than $0.5$ (dotted) using RoSE (red) and TS (green). Colorbars: (a) photons per $58.5 \times 58.5$ nm$^2$; (b) confidence. Scalebar: $500$ nm.

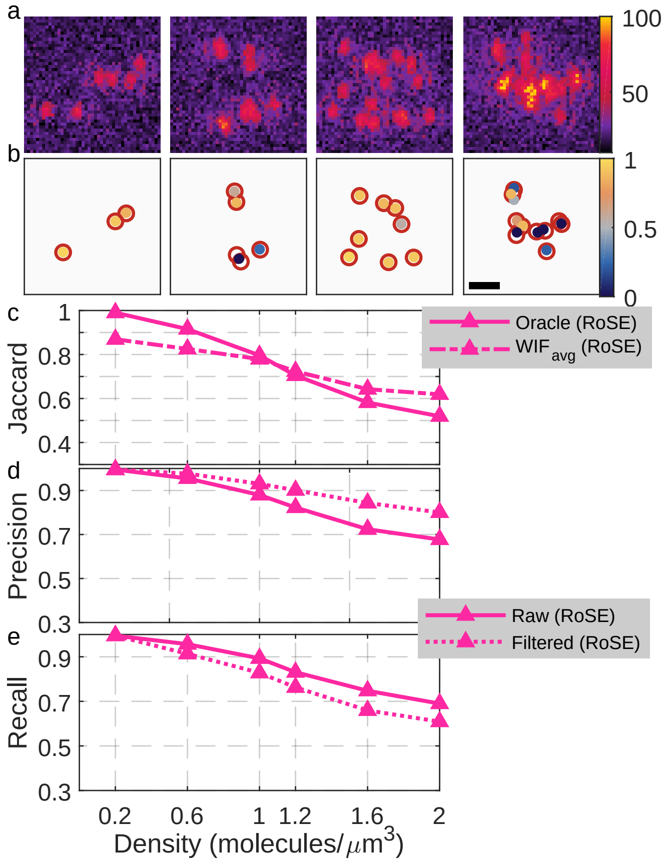

**Fig. S17.** Wasserstein-induced flux $(\mathrm{WIF}_{\mathrm{avg}})$ quantifies 3D localization accuracy without ground truth. (a) From left to right: DH-PSF images of molecules for blinking densities of $0.6$, $1.2$, $1.6$, $2$ mol./$\mu$m$^3$, respectively ($2.5 \times 2.5 \times 0.8$ $\mu$m$^3$ is depth of localization volume). (b) RoSE localizations (colored dots represent calculated confidence) corresponding to images in (a). Open red circles represent ground-truth positions. (c) Jaccard index for RoSE (solid, red) at various blinking densities. The dashed lines represent $\mathrm{WIF}_{\mathrm{avg}}$ for RoSE (red). For each blinking density, 300 independent realizations were used. (d) Precision (detection precision and not localization precision) for all localizations (solid) and localizations with confidence greater than $0.5$ (dotted) using RoSE. (e) Recall for all localizations (solid) and localizations with confidence greater than $0.5$ (dotted) using RoSE. Colorbars: (a) photons per $100 \times 100$ nm$^2$; (b) confidence. Scalebar: $1$ $\mu$m.

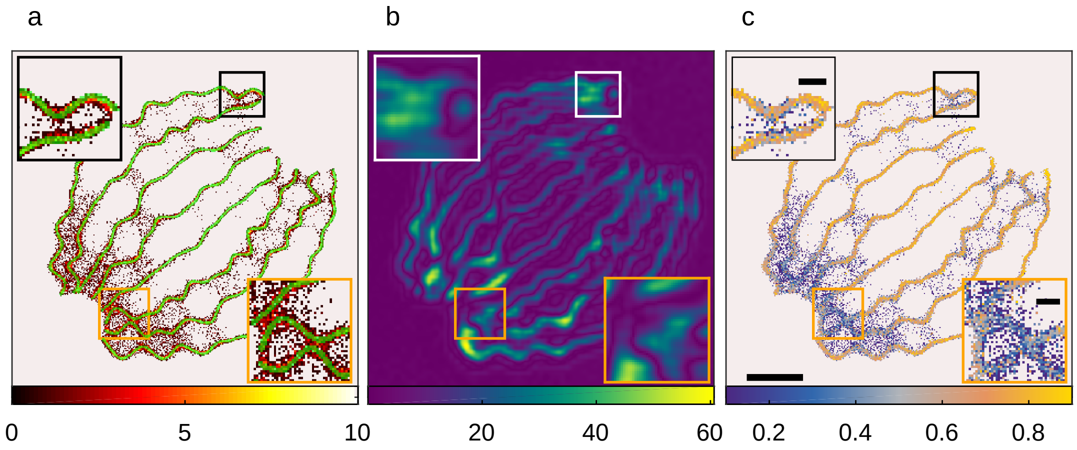

**Fig. S18.** WIF confidence map reveals artifacts in recovering a tubulin network from high-density SMLM data. (a) Recovered structure (red) using FALCON overlaid with the ground truth (green). (b) Error map recovered by SQUIRREL (brighter colors correspond to larger errors). (c) WIF confidence map (brighter colors indicate higher confidence) obtained by averaging localization confidences in each pixel. Colorbars: (a) number of localizations, (b) error, and (c) confidence per $20 \times 20$ nm$^2$. Scalebars: (c) $1$ $\mu$m, insets: $200$ nm

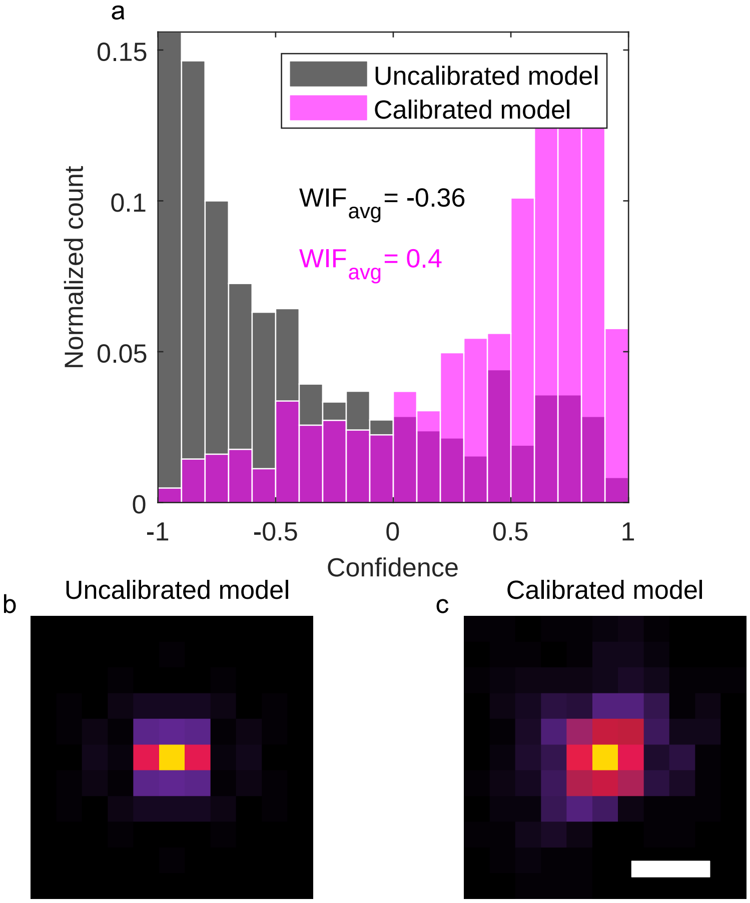

**Fig. S19.** Localization confidences using uncalibrated and calibrated models for the 2D microtubule dataset (Fig. 3). (a) Histograms of WIFs for $600$ isolated images of Alexa Fluor 647 molecules using the uncalibrated ideal PSF model (gray) and the calibrated model (magenta, Materials and Methods). (b,c) PSFs of the uncalibrated and the calibrated model, respectively. Scalebar: (b,c) $500$ nm.

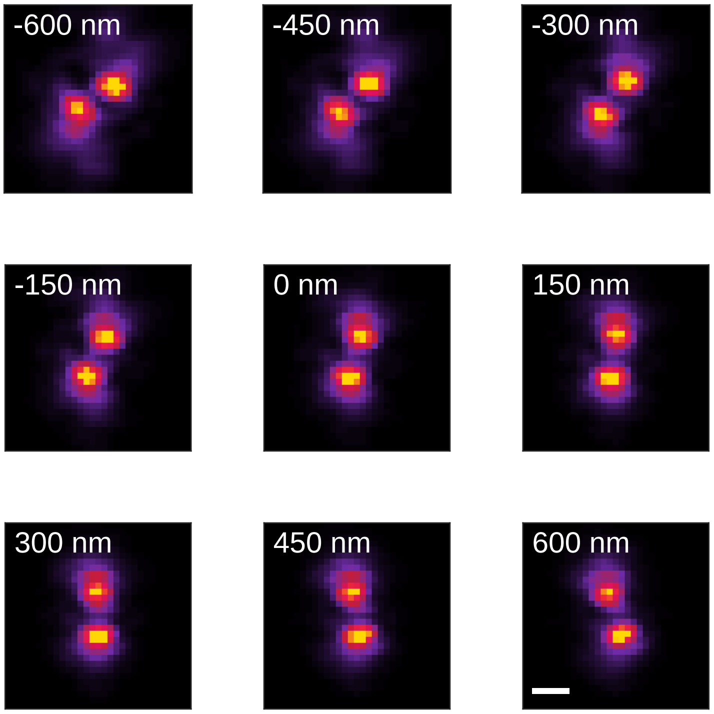

**Fig. S20.** 3D model of the experimental Double-Helix PSF at various axial positions (See Methods in main text for details).

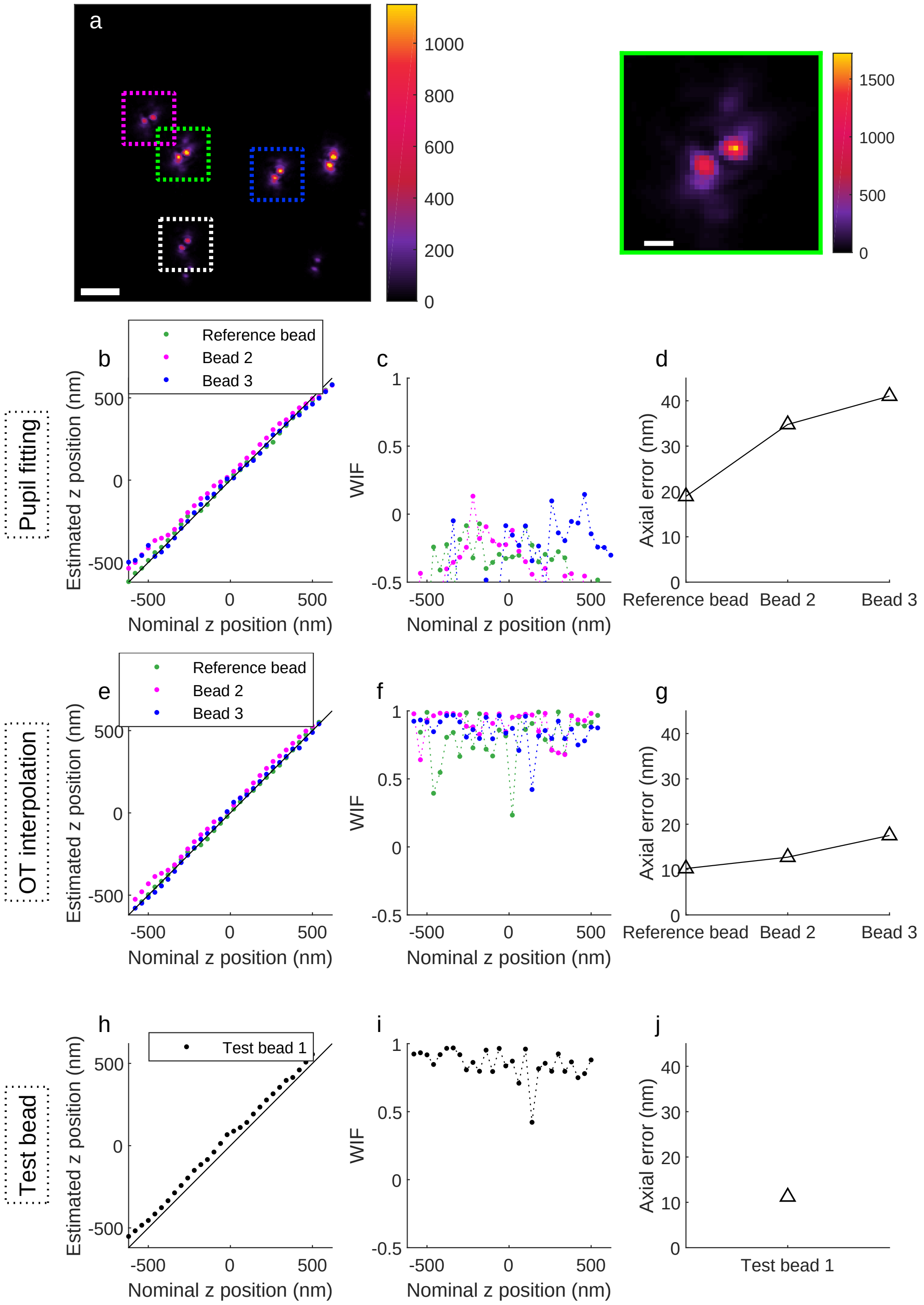

**Fig. S21.** WIF quantifies the accuracy of various 3D PSF models. (a) Palette of fluorescent beads (200 nm diameter) imaged via the DH-PSF with a z-stage step size of $40$ nm within an axial range of $[-620,\ 620]$ nm. The reference bead (green) is chosen such that it has the best fit (obtained using double-Gaussian fitting) to the apparent z positions when moving the microscope stage. Green: reference bead, magenta: bead 2, blue: bead 3, white: test bead. (b) Estimated z positions of beads using RoSE with the PSF model obtained via pupil fitting (the pupil was fitted to the reference bead only). (c) Corresponding WIFs for localizations in (b), computed with same PSF model in (b). (d) Axial error of various beads corresponding to localizations in (b). For each bead, the axial error is equal to the standard deviation of estimated z positions taken at each z-step averaged across all z-steps. (e,f,g) Similar to (b,c,d) but for the PSF model obtained using optimal transport (OT) based interpolation and alignment (See Methods in main text). (h,i,j) Similar to (e,f,g) but for test bead 1 in (a) (enclosed in a white box). Note that test bead 1 is not used when building the PSF model using OT interpolation. Colorbars: (a) detected photons per $160 \times 160$ nm$^2$. Scalebars: (a, left) 5 $\mu$m, (a, right) 1 $\mu$m.

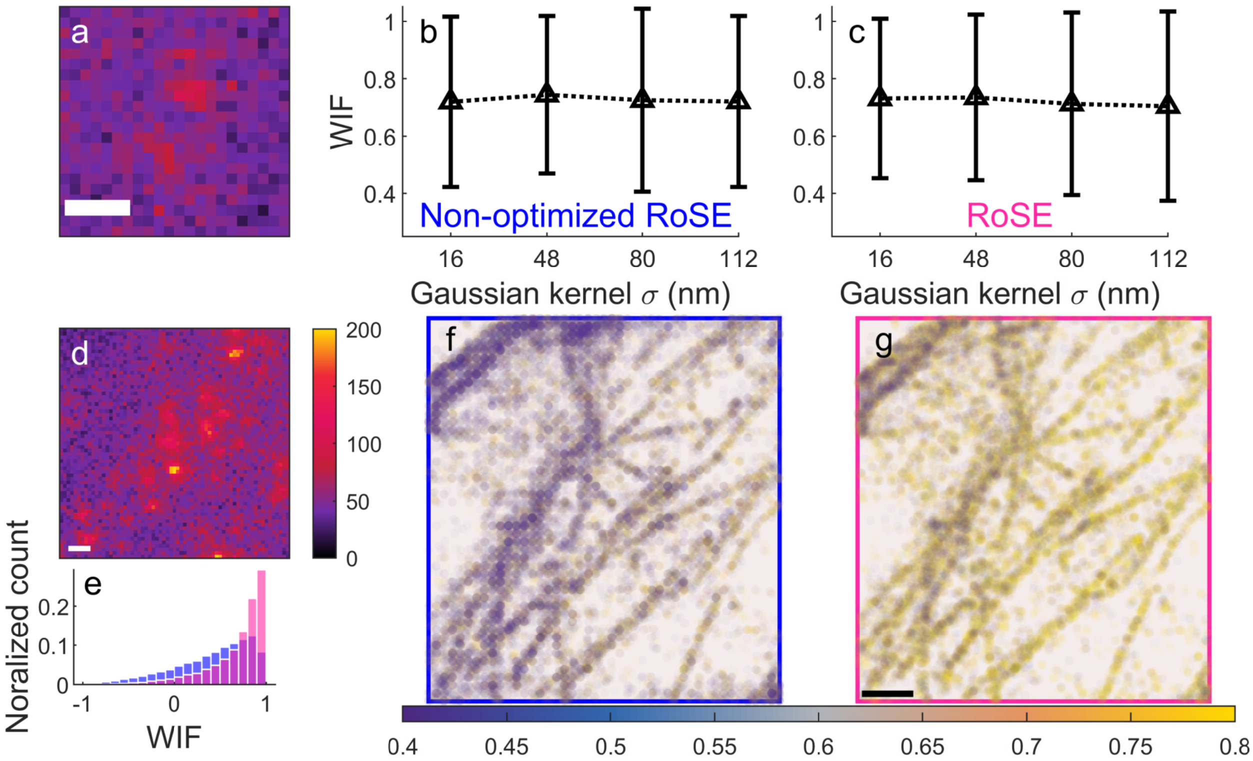

**Fig. S22.** WIF reveals sub-optimal fitting algorithms directly from experimental 3D SMLM data. WIF can be used to quantify the accuracy of RoSE's fitting methods on images of isolated molecules and stochastically activated molecules. (a) Representative DH-PSF image of an isolated SM. (b) Estimated WIFs from 220 isolated SM images (brightness $> 1431$ photons) for non-optimized RoSE (Supplementary Note 4A). The PSF model for SMs was generated by deconvolving a 2D Gaussian kernel of width $\sigma$ from the PSF model obtained from z-stacks of fluorescent beads ($200$ nm diameter). Black triangles indicate mean WIF and solid, and the error bars indicate plus/minus one standard deviation. (c) Similar to (b), but for RoSE. (d) Representative image of stochastically activated SMs of a dense microtubule network (dataset corresponds to Fig. 3). (e) Histograms of confidences or WIFs for non-optimized RoSE (blue) and RoSE (red) corresponding to $10718$ imaging frames across the field of view (FOV) in (d). (f) WIF-density plot of non-optimized RoSE's $36194$ localizations (brightness $> 1000$ photons) corresponding to $10718$ imaging frames across the FOV in (d). Note the gridding artifact present in the localizations. (g) Similar to (f), but for RoSE's $34148$ localizations (brightness $> 1000$ photons). Colorbars: (a,d) detected photons per $160 \times 160$ nm$^2$, (f,g) WIF. Scalebars: (a,d,g) 1 $\mu$m.

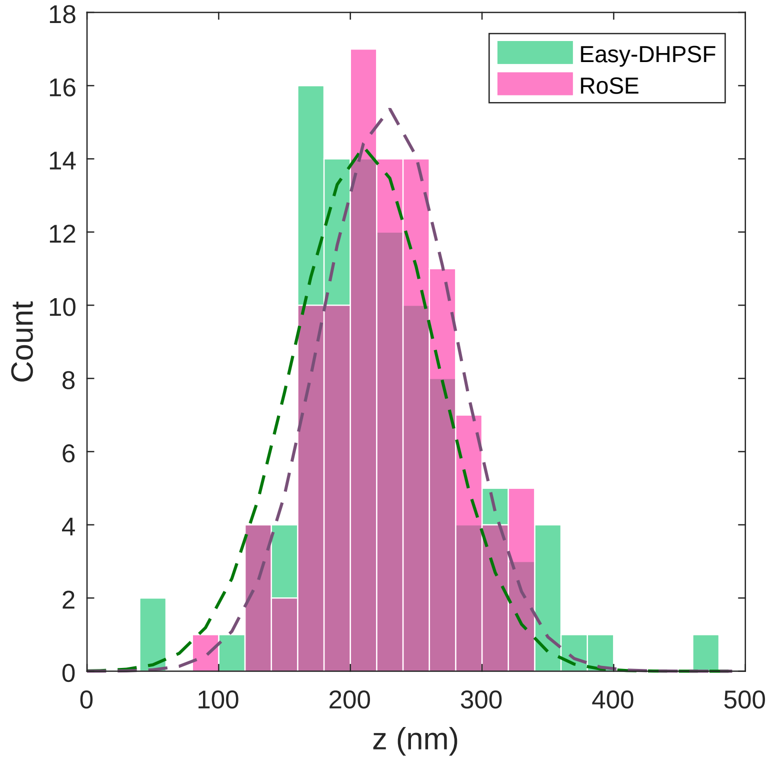

**Fig. S23.** Histograms of the dotted line profile 1 (Fig. 3) along z for Easy-DHPSF (green) and RoSE (red). $95\%$ confidence intervals for the standard deviations of the fitted Gaussian curves were estimated to be $[67, 85]$ nm (Easy-DHPSF) and $[67, 78]$ nm (RoSE).

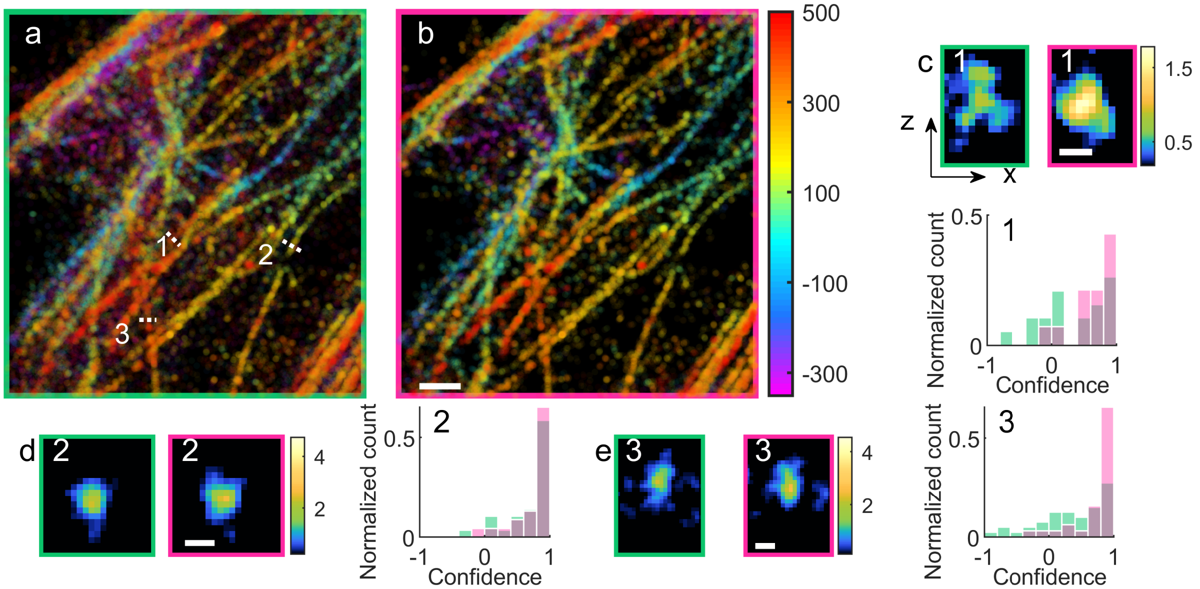

**Fig. S24.** Validation of WIF on 3D experimental images of Alexa Fluor 647-labeled microtubules (dataset corresponds to Fig. 3). (a) 3D SMLM image of Easy-DHPSF localizations with apparent brightness greater than 1300 photons. (b) Similar to (a), but for RoSE's localizations. (c) Transverse (xz) images and WIF distributions of localizations along the dotted line (1) in (a) corresponding to Easy-DHPSF (green) and RoSE (red). Transverse images are obtained by rendering each localization with a 2D, isotropic Gaussian function with a standard deviation of $20$ nm. Mean/median confidence or WIF: $0.36/0.45$ (Easy-DHPSF), $0.62/0.72$ (RoSE). Note the appreciable differences between Easy-DHPSF's reconstruction (green) and that of the RoSE's (red). (d) Similar to (c), but along the dotted line (2) in (a). Mean/median confidence or WIF: $0.68/0.86$ (Easy-DHPSF), $0.73/0.89$ (RoSE). Notice that the high value of WIFs for both algorithms match the higher quality of the reconstructed cross sections. (e) Similar to (d), but along the dotted line (3) in (a). Mean/median confidence or WIF: $0.36/0.49$ (Easy-DHPSF), $0.72/0.85$ (RoSE). The smaller WIFs for Easy-DHPSF are consistent with the observed distortion in the reconstructed cross section. Window widths of the line profiles: (a 1) 716 nm, (a 2) 438 nm, (a 3) 500 nm. Colorbars: (a,b) depth (nm), (c-e) localization density per $20 \times 20$ nm$^2$. Scalebars: (a) 1 $\mu$m, (c-e) 100 nm.

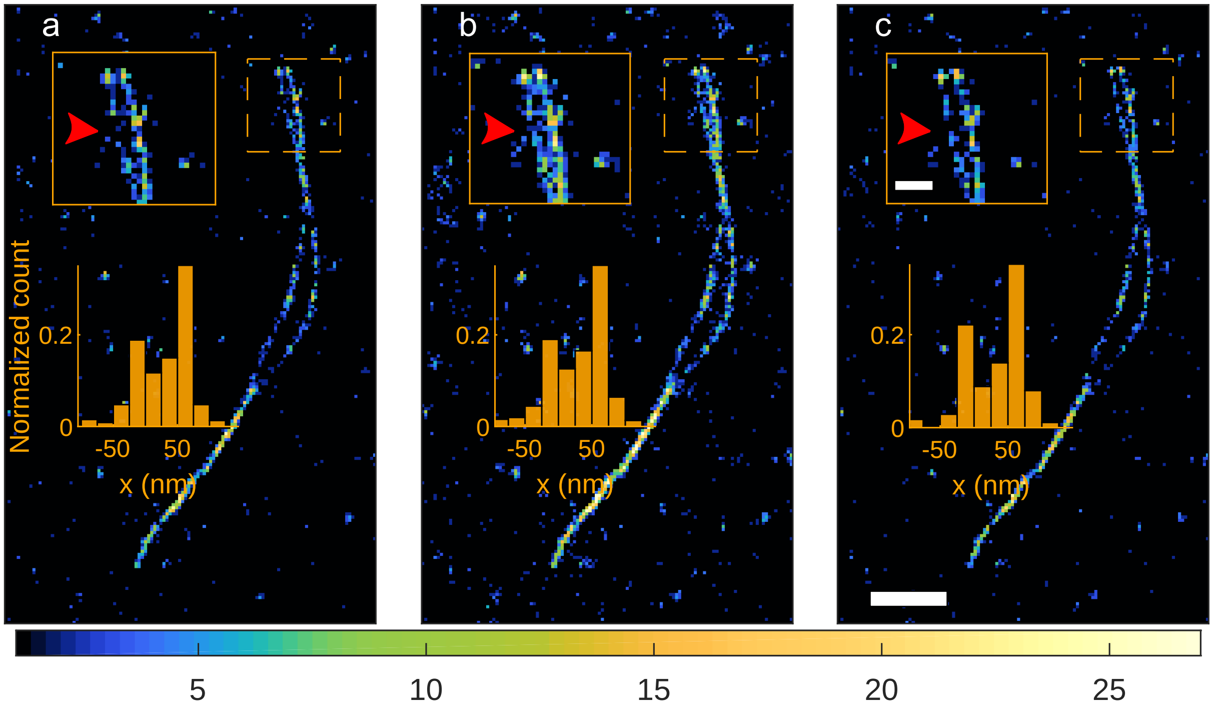

**Fig. S25.** Filtering unreliable localizations via PSF width versus WIF for enhancing reconstruction accuracy. Super-resolution images obtained by TS MLE (Fig. 4b), filtering out localizations (a) with PSF width estimates outside of $[90, 110$ nm$]$, (b) with PSF width estimates outside of $[70, 130$ nm$]$, and (c) with WIF $\leq 0.5$. Insets: Histograms of localizations within the dashed rectangles, projected onto the axis transverse to the fibril, using each filtering strategy (a-c). Colorbar: (a-c) number of localizations per $20 \times 20$ nm$^2$. Scalebar: (a-c) $500$ nm, inset $150$ nm.

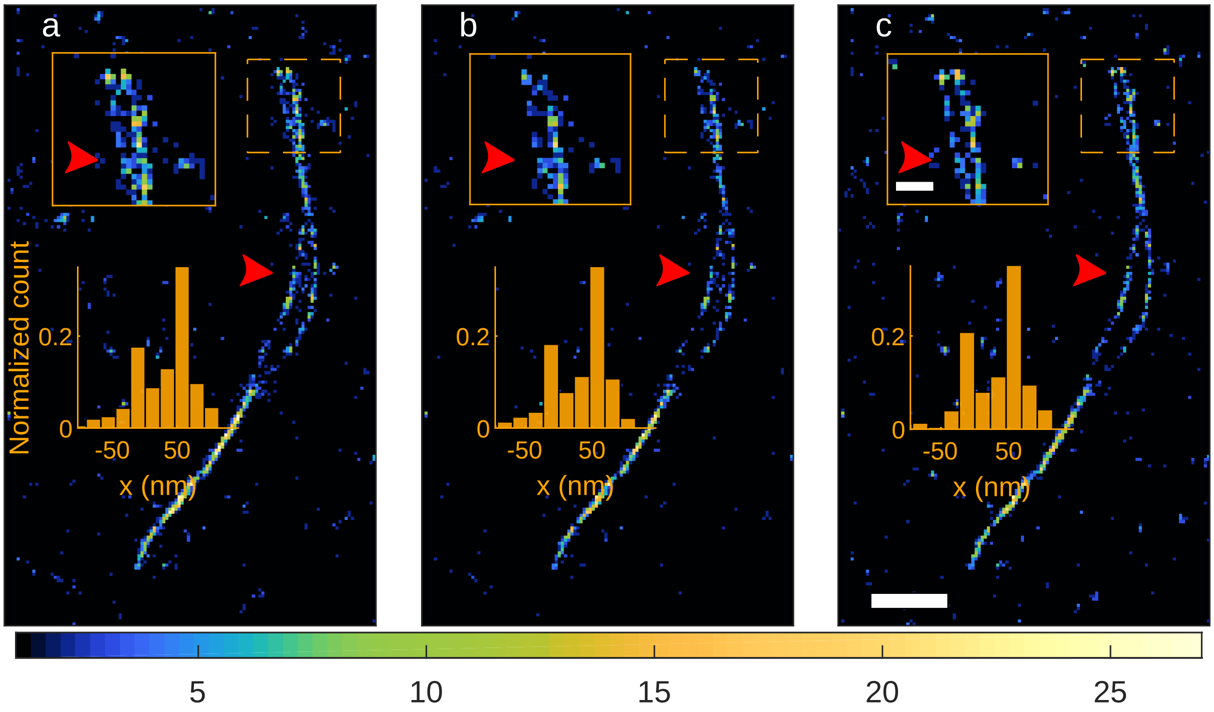

**Fig. S26.** Filtering unreliable localizations using localization precision versus WIF for enhancing reconstruction accuracy. Super-resolution images obtained by TS MLE (Fig. 4b). The images were enhanced by removing localizations (a) with localization precision estimates worse than $6.2$ nm (corresponding to brightness of 300 photons and background of 2.5 per pixel), (b) with localization precision estimates worse than $7.5$ nm (corresponding to brightness of 400 photons and background of 2.5 per pixel), and (c) with WIF $\leq 0.5$. Insets: Histograms of localizations within the dashed rectangles, projected onto the axis transverse to the fibril, using each filtering strategy (a-c). Red arrows indicate regions where bridge artifacts are not filtered by localization precision, but are filtered by WIF. For each molecule, the corresponding localization precision was estimated using Eq. (41). Colorbar: (a-c) number of localizations per $20 \times 20$ nm$^2$. Scalebar: (a-c) $500$ nm, inset $150$ nm.

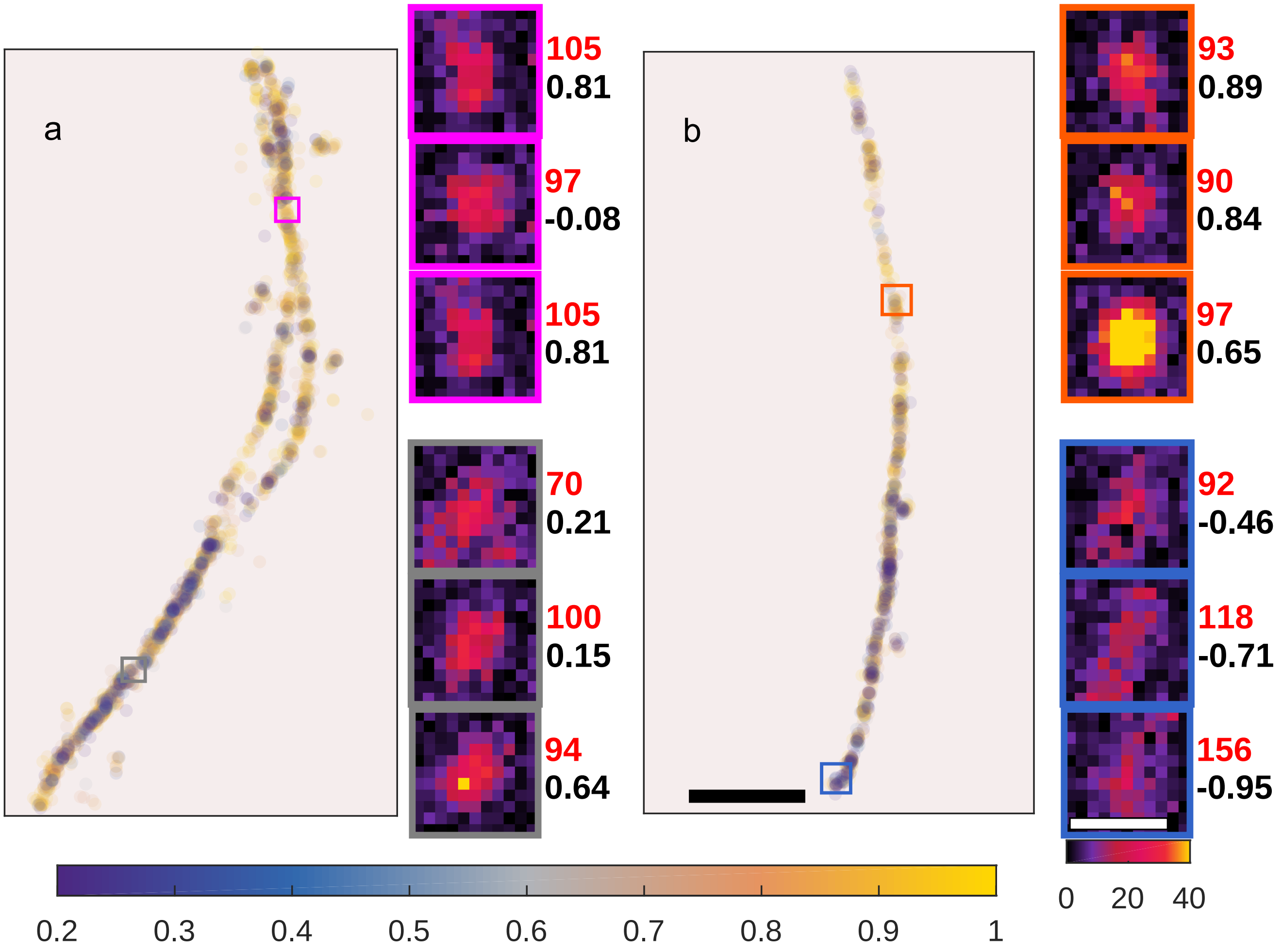

**Fig. S27.** Examples of heterogeneity in Nile red interactions with amyloid fibrils. (a) Density plot of WIFs (with higher-confidence localizations painted last) for bright localizations (>400 photons) on the fibrils. Insets represent example images of Nile red from the colored, boxed regions in (a). Numbers next to insets indicate PSF width estimates (red, nm) and WIF or confidence (black). (b) Similar to (a) but for another fibril. Colorbars: (a,b) confidence, insets: photons/$58.5 \times 58.5$ nm$^2$. Scalebars: (a,b) $500$ nm, inset: $500$ nm.

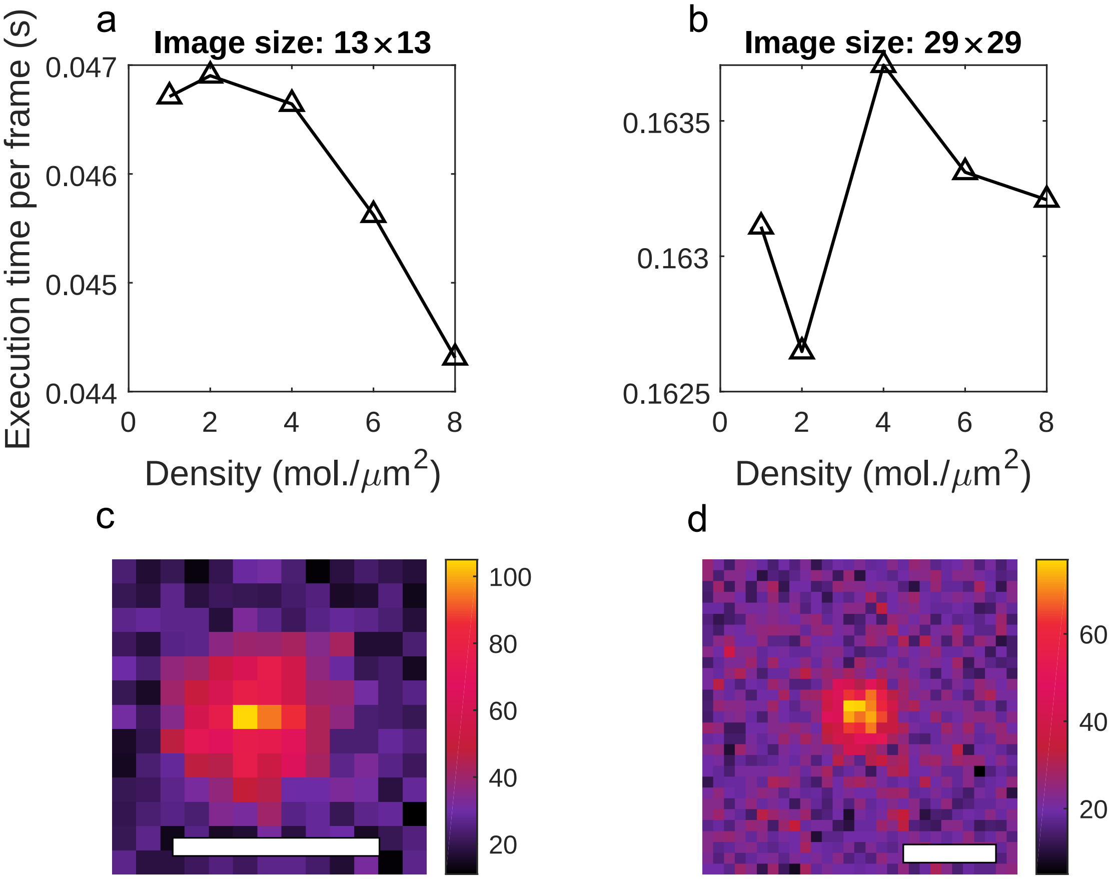

**Fig. S28.** Execution time of computing WIF for various densities and two image sizes. For each density, $200$ independent images of isotropic molecules were analyzed and the background was set to $20$ photons per pixel. Execution time is calculated by averaging the the total time of analyzing $200$ frames. Colorbar: (c,d) photons$/58.5 \times 58.5$ nm$^2$. Scalebar: (c,d) $500$ nm.

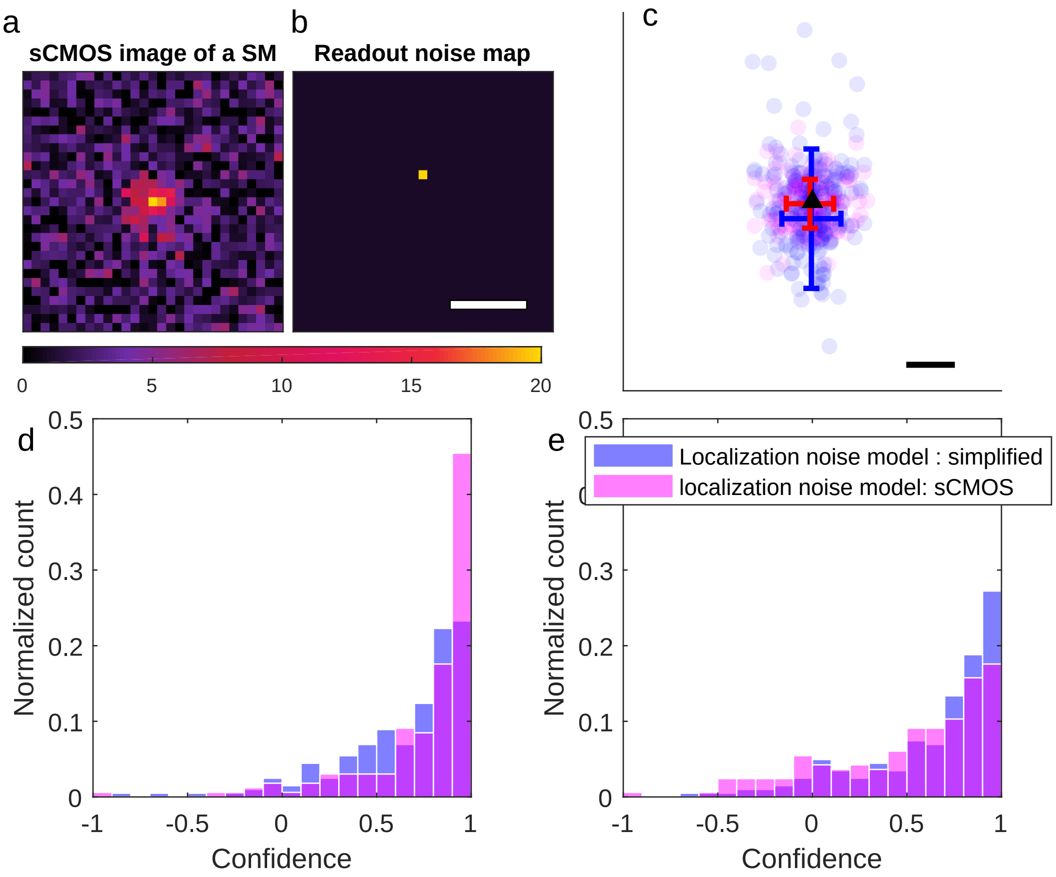

**Fig. S29.** Effect of different measurement noise models on the accuracy of WIF. (a) Representative simulated image of a molecule with a pixel-dependent readout noise modeling the measurement process in sCMOS cameras. (b) Readout noise map (each pixel value represents the standard deviation of the readout Gaussian noise). The readout noise at each pixel was modeled as a Gaussian random variable with mean 0 and standard deviation of 1 photon except for the pixel above the center whose standard deviation was set to 20. The brightness of SM was set to 200 photons with 2 photons per background and a total of 200 realizations were used. (c) Localizations obtained (blue) using a Poisson (simplified) noise model, which ignores the pixel-dependent readout noise, and (magenta) using a shifted Poisson noise model (named sCMOS), which approximates the convolved Poisson and Gaussian distribution with a Poisson one. (d) WIF distributions for localizations in (c) with $\text{WIF}_{\text{avg}} =$ (blue) 0.65 and (magenta) 0.75. Note that the the noise model in WIF algorithm was set to sCMOS for both cases. (e) WIF distributions for localizations in (c) with $\text{WIF}_{\text{avg}} =$ (blue) 0.63 and (magenta) 0.51. Note that the the noise model in the WIF algorithm was set to simplified for both cases. Colorbar: photons per $58.5 \times 58.5$ nm$^2$. Scalebars: (b) 500 m; (c) 20 nm.

 

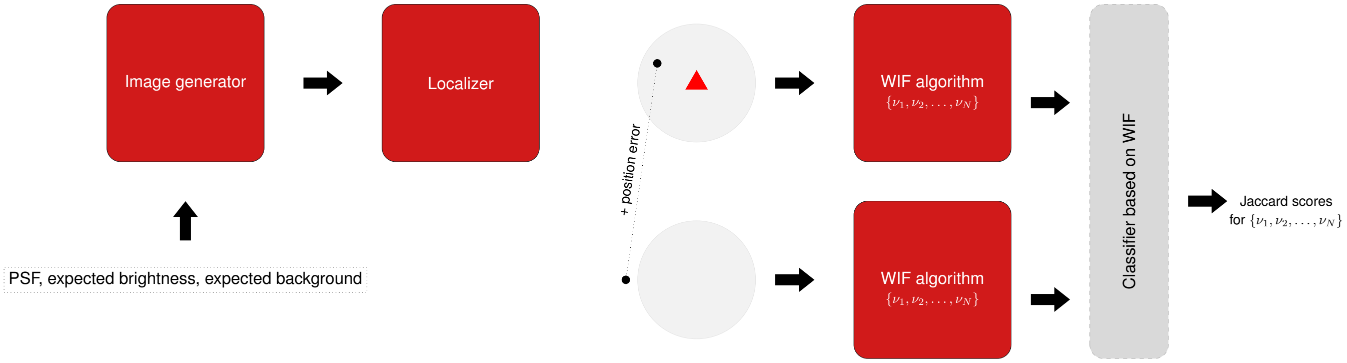

**Fig. S30.** Tuning regularizer strength for optimal performance of WIF. The proposed tuning method accepts a PSF model and basic parameters of the experiment such as expected brightness and background. Next, the method generates images of molecules with various densities. These images are fed into an optimal localization algorithm which outputs accurate localizations. At the same time, these accurate localizations are corrupted by some position errors which can be randomly selected within a certain range. Both accurate and corrupted locaizations are fed into the WIF algorithm for various regularizer strengths. Next, based on a chosen WIF threshold, a classifier assigns a label, accurate or inaccurate, to each localizations. Finally, given the labels, we compute the Jaccard index to quantify the performance of WIF at each regularizer strength and the optimal value of the regularizer is selected as the one which maximizes the Jaccard index.

**Table S2. Conditions and parameters used in computing WIF.**

| Dataset | PSF model | Image pixel size | $2\rho$ | $\nu$ | NA | wavelength |
|---|---|---|---|---|---|---|
| Figs. 1, S3, S4, S6-S8, S10, S11, S15, S16, S29 | ideal standard PSF of isotropic emitter | 58.5 nm | 58.5/2 nm | 0.1 | 1.4 | 637 nm |
| Figs. S12-S14 | ideal standard PSF of isotropic emitter | 58.5 nm | 58.5/2 nm | [0.06,.12] | 1.4 | 637 nm |
| Fig. S18 | ideal standard PSF of isotropic emitter | 100 nm | 100/2 nm | 0.1 | 1.4 | 723 nm |
| Figs. 2, S19 | calibrated linearly-polarized PSF of isotropic emitter | 160 nm | 160/3 nm | 0.1 | 1.4 | 660 nm |
| Figs. 4, 5, S25-S27 | linearly-polarized PSF of isotropic emitter | 58.5 nm | 58.5/2 nm | 0.1 | 1.4 | 610 nm |
| Figs. 3, S21, S22 | experimentally-derived PSF | 160 nm | 160/2 nm | 0.1 | 1.4 | 660 nm |
| Figs. S5, S9, S17 | DH-PSF of isotropic emitter | 100 nm | 100/2 nm | 0.08 | 1.4 | 637 nm |
